# Gut Microbiota Mediates High-Fiber Diet Alleviation of Maternal Obesity-Induced Cognitive and Social Deficits in Offspring

**DOI:** 10.1101/2020.07.16.206714

**Authors:** Xiaoning Liu, Xiang Li, Bing Xia, Xin Jin, Zhenhua Zeng, Shikai Yan, Ling Li, Shufen Yuan, Shancen Zhao, Xiaoshuang Dai, Fei Yin, Enrique Cadenas, Rui Hai Liu, Beita Zhao, Min Hou, Zhigang Liu, Xuebo Liu

## Abstract

Maternal obesity has been reported to be related to the neurodevelopmental disorders in offspring. However, the underlying mechanisms and effective interventions remain unclear. Here, a cross-sectional study on 778 children aged 7-14 years in two cities of China indicates that the maternal obesity is highly associated with the poorer cognition and sociality of their children. Moreover, we also find that the maternal obesity in mice disrupts the behavior and the gut microbiome in the offspring, which are alleviated by a high-fiber diet in either dams or offspring. Co-housing and feces microbiota transplantation experiments reveal a causal relationship between the reshaped microbiota and the behavioral changes. Moreover, treatment of the microbiota-derived short-chain fatty acids exhibits a similar beneficial effect on alleviating the behavioral deficits in offspring. Together, our study purports the microbiota-metabolites-brain axis as a mechanism, and high-dietary fiber intake is a promising intervention against maternal obesity-induced cognitive and social dysfunctions.

## INTRODUCTION

Obesity in women of reproductive age is increasing in prevalence worldwide due to intake of high-calorie, low-fiber diets, consumption of manufactured foods with high in sugar, and sedentary lifestyle (Hanson et al., 2017; Poston et al., 2016). Maternal obesity before and during pregnancy is widely recognized to have short-term and long-term adverse health outcomes for both the mothers and their children. Evidences from both animal and human studies implicate that maternal obesity induced by dietary intervention leads to obesity, diabetes, increased blood pressure, and even behavioral changes in the offspring (Drake and Reynolds, 2010; Godfrey et al., 2017; Harris et al., 2020), which indicates maternal obesity may be one of the influences contributing to the “developmental origins of health and disease” (Drake and Reynolds, 2010). The high prevalence of maternal obesity means that the determination of any such long-term effects is now an urgent priority (Heslehurst et al.). Recent UK and US nationally distributed longitudinal studies has shown increased risks for poorer cognitive outcomes and autism spectrum disorders (ASD) in children of mothers with obesity before pregnancy (Basatemur et al., 2013; Pugh et al., 2015). Work in rodents also showed that maternal high-fat diet (MHFD) induces long-term cognitive deficits across several generations recently (Sarker and Peleg-Raibstein, 2018). Despite the potential public health importance, few cohort studies have been done to examine the neurobiological mechanism by which maternal obesity affects offspring behavior and brain function.

A non-genetic, yet heritable contributor to behaviors may be the microbiota, a community of microorganisms harbored in gastrointestinal (GI) tract that impacts development and function of the immune, metabolic, and nervous systems (Cho and Blaser; Moeller et al., 2018; Santiago et al.). The intestinal ecosystem is thought to be established at or soon after birth, facilitated by vertical transmission and exposure to and/or ingestion of environmental flora (Ferretti et al., 2018; Moran et al., 2018). Thus, maternal influences on the offspring’s microbiome are significant, and may potentially alter the risk of mental impairment. Indeed, maternal obesity has been associated with alterations in the gut microbiome in offspring in both human and non-human primates (Batterham et al., 2002; Chu et al., 2016). A recent study on *Macaca fuscata* (Japanese macaque) has reported that a high-fat maternal diet, but not obesity *per se*, re-structured the offspring’s intestinal microbiome. Specifically, maternal obesity negatively impacts a subset of bacteria in the offspring gut and selective re-introduction of *Lactobacillus (L.) reuteri* restores social deficits in offspring (Buffington et al., 2016). A recent study also identifies differences between Alzheimer’s disease and normal controls in 11 genera from the feces and 11 genera from the blood (Li et al., 2019). Given emerging reports that link gut microbiota to brain function, it has been speculated that changes in the gut microbiome may also be relevant to cognitive impairments in offspring induced by maternal obesity.

Dietary fiber as the seventh nutrient is widely reported to regulate the gut microbiome, providing an important substrate to the community of microbes (microbiota) that inhabits the distal gut (Sonnenburg and Sonnenburg, 2014). Unlike humans, who produce ∼17 gastrointestinal enzymes to digest mostly starch, the gut microbiota produces thousands of complementary enzymes with diverse specificities, enabling them to depolymerize and ferment dietary polysaccharides into host-absorbable – short chain fatty acids (SCFAs) (Kaoutari et al., 2013). SCFAs have various effects on the host brain. For example, acetate crosses the blood-brain barrier, where it is taken up and regulates the activity of hypothalamic orexigenic neurons (Farhadi et al.). Notably, SCFAs are key molecules that modulate microglia maturation, morphology and function (Erny et al., 2015). Additional evidences suggest that dietary manipulation of the maternal microbiome in pregnancy with fiber has beneficial effects for both offspring immune function and metabolism (Kimura et al., 2020; Thorburn et al., 2014). However, it still is unknown whether dietary fiber restores cognitive and social behavioral deficits in offspring from obesity mothers.

Here we report that a) maternal obesity induces cognitive and social behavioral deficits in human and mouse offspring; b) maternal high fiber intake improves offspring social behavioral deficits induced by maternal high fat diet (MHFD) by restructuring the gut microbiome transmitted from mother to offspring; c) mother-to-offspring gut microbiome mediates maternal high fat diet (MHFD) induced learning and memory impairments in offspring; d) offspring high fiber intake improves gut microbiome, aberrant spliceosome alterations and restores HFD-induced cognitive and social behavioral deficits in MHFD-CD offspring; e) oral treatment with a mixture of acetate and propionate reverses behavioral deficits in MHFD-CD Offspring.

## RESULTS

### Maternal Prepregnancy Overweight and Obesity Are Associated with Impaired Child Neurodevelopment

To examine the association between maternal pregnancy weight and neurodevelopment in offspring, 778 children (403 boys and 375 girls) were eligible for follow-up at 7-14 years of age, whose parents completed the social competence scale of the Child Behavior Checklist, which included 20 social competence items with three social competence subscales measure competencies: activities (e.g., sports and hobbies); social subjects(e.g., friendships and interpersonal skills); and school performance (e.g., performance, learning ability and school problems). Demographics of the study population are shown in **Table 1**. Compared with normal weight mothers, overweight & obese mothers had the highest proportions of overweight & obese children (*p* < 0.05). Additionally, overweight & obese mothers had on average lower educational attainment and lower family income. On average, all underweight, normal and overweight & obese mothers gained pregnant weight gain as recommended. Compared with children of normal weight mothers, children of overweight & obese mothers had significantly lower score of social competence (*p* < 0.01) **(Table 1; Figure S1A)**. Consistent with other countries cohort studies of poorer cognitive performance and increased risk of autism spectrum disorders in children of obese mothers, children of Chinese overweight & obese mothers had significantly reduced scores of social subjects and school performance, but not activities, indicating lowered social and learning ability (*p* < 0.05) **(Table 1; Figures S1B-S1D)**. Further analysis revealed that significantly reduced social competence were more pronounced in boys than in girls, which could be due to lower school performance scores (*p* < 0.01) **(Figures S1A and S1B)**. Taken together, these data indicate that maternal obesity could be more likely to cause cognitive and social impairments in their male offspring.

**Table 1.** Participant Characteristics and Social Competence among Children.

| Characteristics | Maternal Pre-pregnancy BMI |  |  | P value |
| --- | --- | --- | --- | --- |
|  | Underweight<br>(BMI<18.5)<br>N=132 | Normal<br>(BMI 18.5-23.9)<br>N=533 | Overweight<br>& Obesity<br>(BMI >23.9)<br>N=79 |  |
| Child's age (years)<br>(Mean ± SD) <sup>a</sup> | 10.32±2.01 | 10.39±1.86 | 10.42±1.95 | 0.955 |
| Child's gender <sup>b</sup> |  |  |  |  |
| Male, % | 52.3 | 47.3 | 49.4 | 0.941 |
| Child's BMI <sup>b</sup> |  |  |  | 0.012 |
| Underweight, % | 15.4 | 11.6 | 15.9 |  |
| Normal, % | 73.2 | 69.1 | 53.6 |  |
| Overweight & Obesity, % | 11.4 | 19.4 | 30.4<sup>+</sup> |  |
| Breast feeding <sup>a</sup> | 81.8 | 82.7 | 79.7 | 0.801 |
| Maternal Education level <sup>b</sup> |  |  |  | 0.001 |
| Secondary high school or lower, % | 8.3 | 14.8 | 36.7 |  |
| High school, % | 30.3 | 19.9 | 15.2 |  |
| Diploma in higher education or Bachelor's degree, % | 59.1 | 63.8 | 48.1 |  |
| Master's degree or higher, % | 2.3 | 1.5 | 0.0 |  |
| Maternal pregnant weight gain (Mean ± SD) <sup>a</sup> | 15.70±5.97 | 14.24±5.51 | 11.57±6.10 | 0.000 |
| Family income (RMB) <sup>b</sup> |  |  |  | 0.000 |
| < 80,000, % | 5.3 | 9.6 | 27.8 |  |
| 80,000–150,000, % | 33.3 | 32.5 | 32.9 |  |
| 150,000–300,000, % | 39.4 | 39.0 | 30.4 |  |
| > 300,000, % | 21.9 | 19.0 | 8.9 |  |
| Children's social competence (Mean ± SD) <sup>a</sup> | | | | |
| Total | 17.05±4.06 | 17.10±3.72 | 15.65±3.79<sup>*</sup> | 0.005 |
| Ability | 4.89±2.36 | 4.90±2.18 | 4.36±2.21 | 0.147 |
| Social | 6.80±2.00 | 6.78±1.87 | 6.16±1.92<sup>*</sup> | 0.045 |
| School | 5.36±0.63 | 5.43±0.67 | 5.13±0.95<sup>*</sup> | 0.013 |

<sup>a</sup> Kruskal-Wallis test;<sup>b</sup> $\chi^2$ test;<sup>+</sup> Significant difference from underweight maternal BMI category with significance level at

### Maternal Obesity Induces Cognitive and Social Behavioral Deficits in Mouse Offspring

In human studies, confirmation of causation and identification of mechanisms linking maternal obesity with offspring neurodevelopment are difficult. To investigate how maternal obesity affects offspring neurodevelopment, female C57BL/6 mice were fed either control diet (CD) or high-fat diet (HFD) for 12 weeks. As expected, MHFD significantly increased maternal weight **(Figure S2A)**. Females were then paired with males to produce offspring that were fed with a control diet after weaning (3 weeks) (**Figure 1A**). There was no significant difference in offspring weight between maternal diet groups at 8–10 weeks old or at 7 months old (**Figure S2B**), at which the behavioral experiments were performed. There were no sexual differences between male and female offspring in the behavior test. At 8-10 weeks old, to assess working memory and long-term memory, Y-maze test and novel object recognition test were performed, respectively. Compared with MCD-CD offspring, MHFD-CD offspring had fewer spontaneous alternation and showed a discrimination index between the novel and familiar object close to −0.2, indicating impairments of working memory and long-term memory (**Figures 1B, 1C, S2C and S2D**). Consistent with a previous report, MHFD-CD offspring had impaired sociability and showed no preference for social novelty by a three-chamber sociability test (*p* < 0.01) (**Figures 1D and 1E**). Furthermore, MHFD-CD mice still showed significant cognitive impairment but not social, compared with MCD-CD offspring at 7 months of age (**Figures 1B-1E**) (*p* < 0.05). Taken together, these data indicate that maternal obesity leads to offspring’s memory and social deficits.

**Figure 1.**
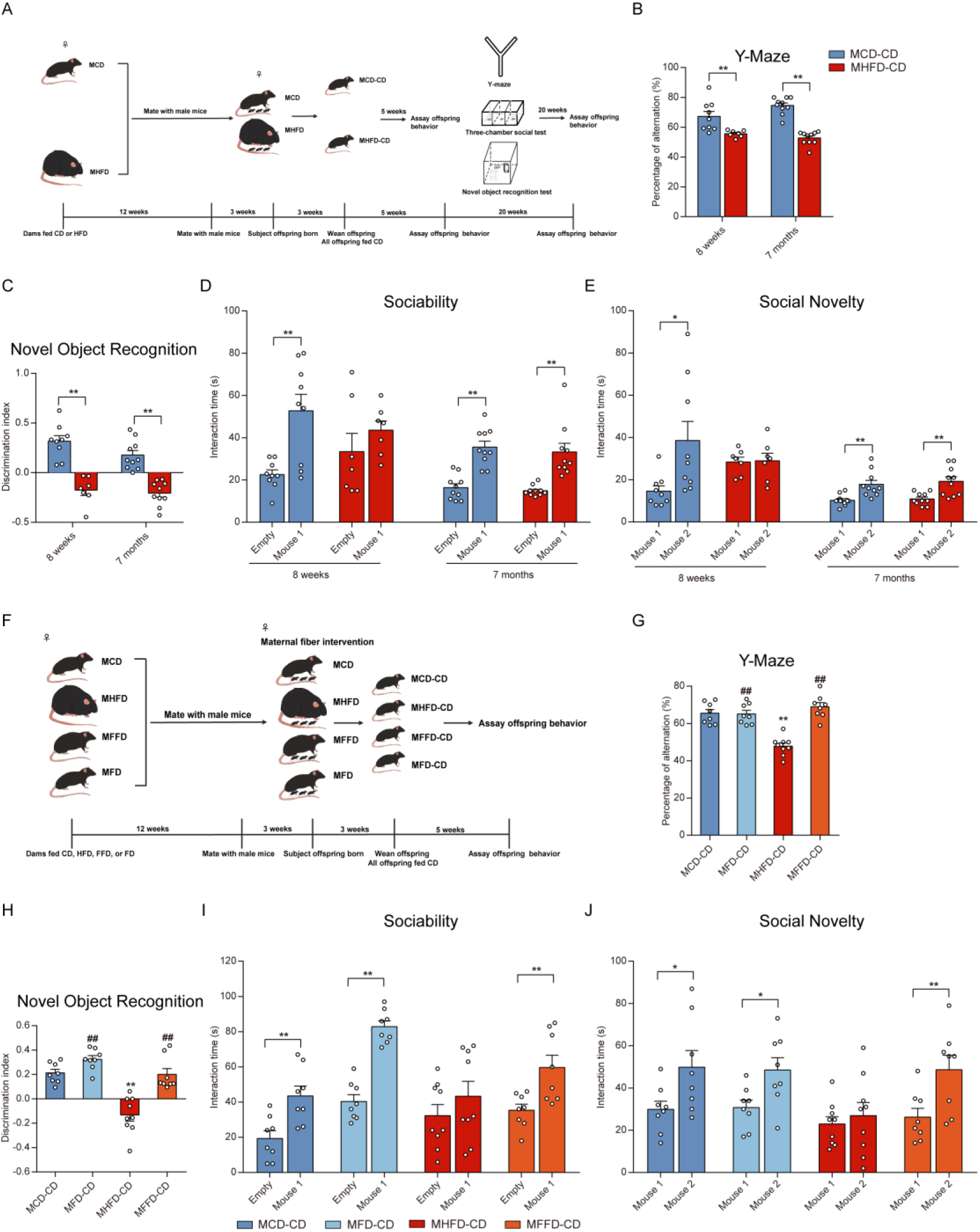
Maternal Obesity and Dietary Fiber Intake Imprint the Cognitive and Social behaviors of Mouse Offspring. (A) Schematic of the maternal diet regimen and breeding. (B) For Y-maze, spontaneous alternations were recorded (n=7-10 mice/group). (C) For the novel object recognition test, the discrimination index between the novel and familiar object were recorded (n=7-10 mice/group). (D) In the sociability test, the time spent on interacting with a mouse or with an empty wire cage were recorded (n=7-10 mice/group). (E) In the social novelty test, the time spent on interacting with a novel versus a familiar mouse (n=7-10 mice/group). (F) Schematic of the maternal dietary fiber supplementation. (G-J) Behavioral phenotypes in offspring after maternal dietary fiber supplementation, including (G) Y-maze, (H) the novel object recognition test, (I) sociability, and (J) preference for social novelty (n = 8-9 mice/group). Data presented as mean ± SEM. Statistical analyses were performed by one-way ANOVA with Tukey’s multiple comparison test (for B, C, G and H) or by paired two-tailed Student’s t test (for D, E, I, and J). For Tukey’s multiple comparison test, \**p* < 0.05, \*\**p* < 0.01, compared with the MCD-CD group, ^#^*p* < 0.05, ^##^*p* < 0.01, compared with the MHFD-CD group. For Student’s t test, \**p* < 0.05, \*\**p* < 0.01. See also Figure S2.

### High Fiber Intake in Maternal Diet Restores Maternal Obesity-Induced Cognitive and Social Deficits in Offspring

To investigate whether dietary fiber alleviates maternal obesity-induced offspring neurodevelopment dysfunction, female mice were fed with one of four diets: control diet (MCD), high-fat diet (MHFD), high-fat/high-fiber diet (a high-fat diet with inulin as a source of fiber, MFFD), and a control diet with inulin (MFD) for 12 weeks. Females were then paired with males to produce offspring that were fed with a control diet after weaning (**Figure 1F**). Consistent with reports of lower fertility in obese mothers (Poston et al., 2016), the litter size was reduced (**Figure S2G**) and latency to first litter increased in female mice fed HFD (**Figure S2H**). Relative to the ‘‘standard’’ HFD, a diet comprised of dietary fiber reduced weight gain and markedly increased fertility (*p* < 0.01) (**Figures S2E-S2H**). There was no significant difference in offspring weight between maternal diet cohorts at 8–10 weeks old (**Figure S2I**). Fecal samples were collected and behaviors in offspring were assessed when mice were 8-10 weeks old. Strikingly, maternal dietary fiber intake significantly improved working memory and long-term memory (*p* < 0.01), as well as sociability and the preference for social novelty in MHFD-FD offspring (*p* < 0.05) (**Figures 1G-1J, S2J-S2N**). Taken together, these data indicate that maternal dietary fiber intake could protect against maternal obesity-induced cognitive and social behavioral deficits in offspring.

### High Fiber Intake in Maternal Diet Restores Maternal Obesity-Induced Synaptic Impairment in Offspring

The ultrastructures of synapses in the hippocampus were examined. An analysis of postsynaptic density (PSD) ultrastructure revealed that the length and width of PSD were significantly elevated in the MFFD-CD group, compared with the MHFD-CD group (*p* < 0.01) (**Figures 2A and 2B**). In line with PSD ultrastructural alterations, the expressions of *PSD-95*, a major component of PSD, was also restored in MFFD-CD offspring (**Figures 2C and S3A**), compared with MHFD-CD offspring as assessed by qPCR and immunofluorescence. Accordingly, the mRNA levels of synapse-related *FXR1*, *GluA2* and *BDNF* were decreased in the offspring derived from the HFD-treated dams, but not in the MFFD-CD offspring (**Figures 2C** and **S3B**).

**Figure 2.**
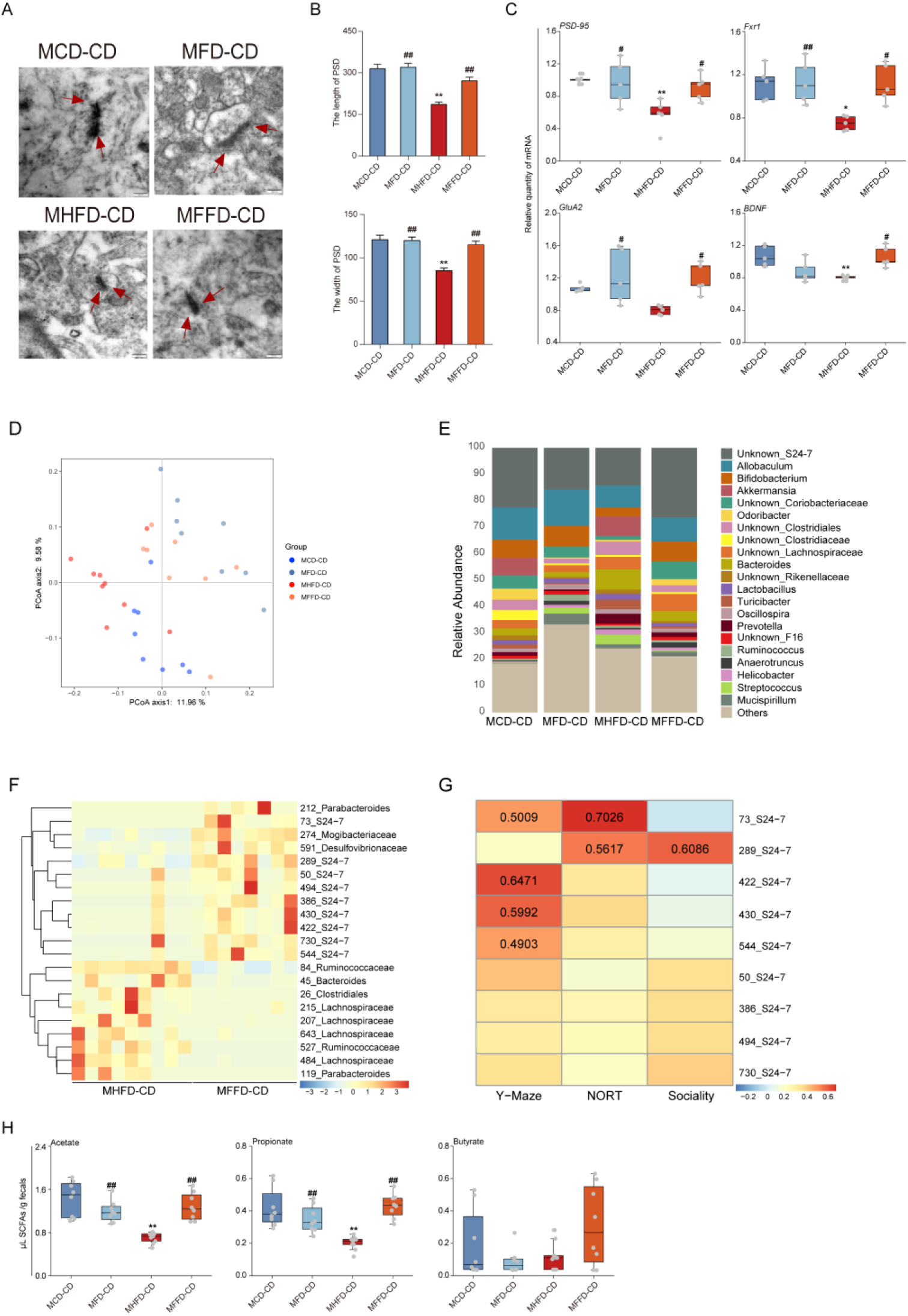
High Fiber Intake in Maternal Diet Restores Synaptic Impairment and Re-shapes the Gut Microbiome of Offspring. (A) Representative images of ultrastructure of synapse. (B) The length (up) and width of PSD (down) (n = 3 mice/group). (C) The mRNA expressions of *PSD-95*, *Fxr1*, *GluA2*, and *BDNF* in the hippocampus (n = 5 mice/group). (D) Principal coordinates analysis (PCoA) of unweighted UniFrac distances from the averaged rarefied 16S rRNA gene dataset (*p* = 0.0001, R^2^ = 0.1993) (n = 8-9 mice/group). (E) The relative abundance of bacteria at the genus level. All genera with an average relative abundance below 1% were grouped to “others”. (F) A Z-score scaled heatmap of different OTUs identified by Wilcoxon rank-sum test between MHFD-CD and MFFD-CD with *p* ≤ 0.01. (G) Abundance of selected taxas in the offspring microbiome is correlated with behavior of offspring. Spearman’s rank correlation between the microbiome and mouse behavior. If *p* value<0.05 for significant correlations, R is noted. NORT: object recognition test (H) The concentrations of short-chain fatty acid (SCFA) including acetate, propionate and butyrate in offspring feces (n = 8-9 mice/group). Data of (B) and (C) presented as mean ± SEM. Data of (H) presented as median ± interquartile range. Statistical analyses were performed by one-way ANOVA with Tukey’s multiple comparison test (for B, C and H). \**p* < 0.05, \*\**p* < 0.01, compared with the MCD-CD group, ^#^*p* < 0.05, ^##^*p* < 0.01, compared with the MHFD-CD group. See also Figure S3 and Figure S4.

Social behaviors are mediated by the prefrontal cortex (PFC). Given that microglia dysregulation was reported in a range of neuropsychiatric conditions including ASD, the expression levels of microglia surface molecule *CD31*, *CSF1R*, and *F4/80*, as well as *Mafb*, an important transcription factor of the adult microglia program (Matcovitch-Natan et al., 2016), were assessed. Maternal obesity lowered *Mafb* mRNA level and increased *CD31* mRNA level in MHFD-CD offspring, which were restored by enrichment of the diet with dietary fiber (**Figure S3C**). Furthermore, the microglia-neuron interaction dysfunction, such as reductions in the levels of *CX3CL1*, *NGF* and *DAP12*, in MHFD-CD offspring were ameliorated in MFFD-CD offspring (**Figure S3D**).

Likewise, unlike the MHFD-CD offspring, the expression of *FXR1*, *FXR2*, *PSD-95* and *Tdp2* were increased in MFFD-CD offspring (**Figures S3E andS3F**). These findings were consistent with reports of shared mechanisms underlying both social and cognitive impairments (State and W). Taken together, these data indicate that maternal high fiber intake protects offspring against synaptic impairment and disruption of microglia maturation in hippocampus and PFC, which were induced by maternal obesity.

### High Fiber Intake in Maternal Diet Re-shapes the Gut Microbiome in Both Mother and Offspring Mice

Maternal obesity has been shown to alter the gut microbiome of offspring (Chu et al., 2016; Ma et al., 2014; Steegenga et al., 2017) and microbial reconstitution could reverse social deficits (Buffington et al., 2016). Dietary fiber is a well-recognized prebiotic known to promote growth of select beneficial bacteria in colon (Gibson, 1999). Hence, we next examined whether dietary fiber’s protection against HFD-induced behavioral deficits would correlate with microbiota alterations. Analysis of microbiota composition by 16S sequencing indicated that there was a remarkable change in the maternal microbiome composition but not community richness between MHFD group and MFFD group **(Figures S4A and S4B)**. Maternal dietary fiber intake also corrected some of the HFD-induced changes in microbiota that were observed at the phylum level and order level **(Figures S4C and S4D)**. Specifically, S24-7 (OUT_73_S24-7, OTU_127_S24-7, OUT_187_S24-7 et al.), *Bifidobacterium animalis* (424_Bifidobacterium_animalis), *Prevotella* (OUT_21_Prevotella) and Clostridiales (OTU_276_Clostridiales and OTU_435_Clostridiales and OTU_277_Clostridiales) were prevalent in most MFFD samples and absent from most or all MHFD samples (**Figure S4E**).

While bacterial α-diversity did not differ between the offspring from any diet group (**Figures S4F**), the unweighted Unifrac distance of the MFFD-CD mice was different from MHFD-CD mice, similar to that in observed in their dams, which revealed that dietary fiber regimen altered the microbiota composition (**Figures 2D and S4A**). Differentiating bacterial taxa in offspring belong predominantly to the Clostridiales and Bacteroidales orders, with single representatives from the Actinobacteria, Firmicutes and Tenericutes phyla (**Figures 2E, S4G and S4H**). Specifically, 21 differentiating bacterial taxa were identified between MHFD-CD and MFFD-CD by Wilcoxon rank-sum test analyzing OTUs, nine of which belonged to S24_7 family (**Figure 2F**). The abundance of five bacterial OTUs positively correlated with increased cognitive and social behaviors in mice using Spearman’s rank correlation (**Figure 2G**) (R> 0.4, *p* < 0.05). The association of specific bacterial species with MFFD-CD samples, which are also highly correlated with increased cognitive- or social-relevant behaviors, supports the hypothesis that specific bacteria may contribute to the protection of dietary fiber against cognitive and social deficits in offspring, induced by maternal high fat diet.

Microbial metabolites in the gut could impact neurological outcomes (Wang et al., 2018). SCFAs in offspring feces were next examined, which are derived from microbial fermentation of dietary fibers and are likely to have broad impacts on various aspects of host physiology. Dietary fiber treatment marked improved MHFD-induced decreases in the feces levels of acetate, propionate, but not butyrate in offspring (**Figure 2H**) (*p* < 0.05).

### Gut Microbiota Mediates Maternal Obesity-Induced Cognitive and Social Deficits in Offspring

The maternal gut microbiome can be vertically transmitted to their offspring by breastfeeding and lead to longer-lasting colonization of the offspring gut (Ferretti et al., 2018; Moran et al., 2018). To investigate whether gut microbiota plays an important role in MHFD-induced behavioral abnormalities, the fecal microbiota from female MHFD and MFFD mice were transplanted into antibiotics-treated adult female mice, respectively (**Figures 3A, S5A and S5B**). Offspring from the dams inheriting MHFD donor microbiota did show impaired memory and social behaviors (**Figures 3B-3E, S5C and S5D**). By contrast, offspring from the dams inheriting MFFD donor microbiota had no behavioral abnormalities (**Figures 3B-3E, S5C and S5D**).

**Figure 3.**
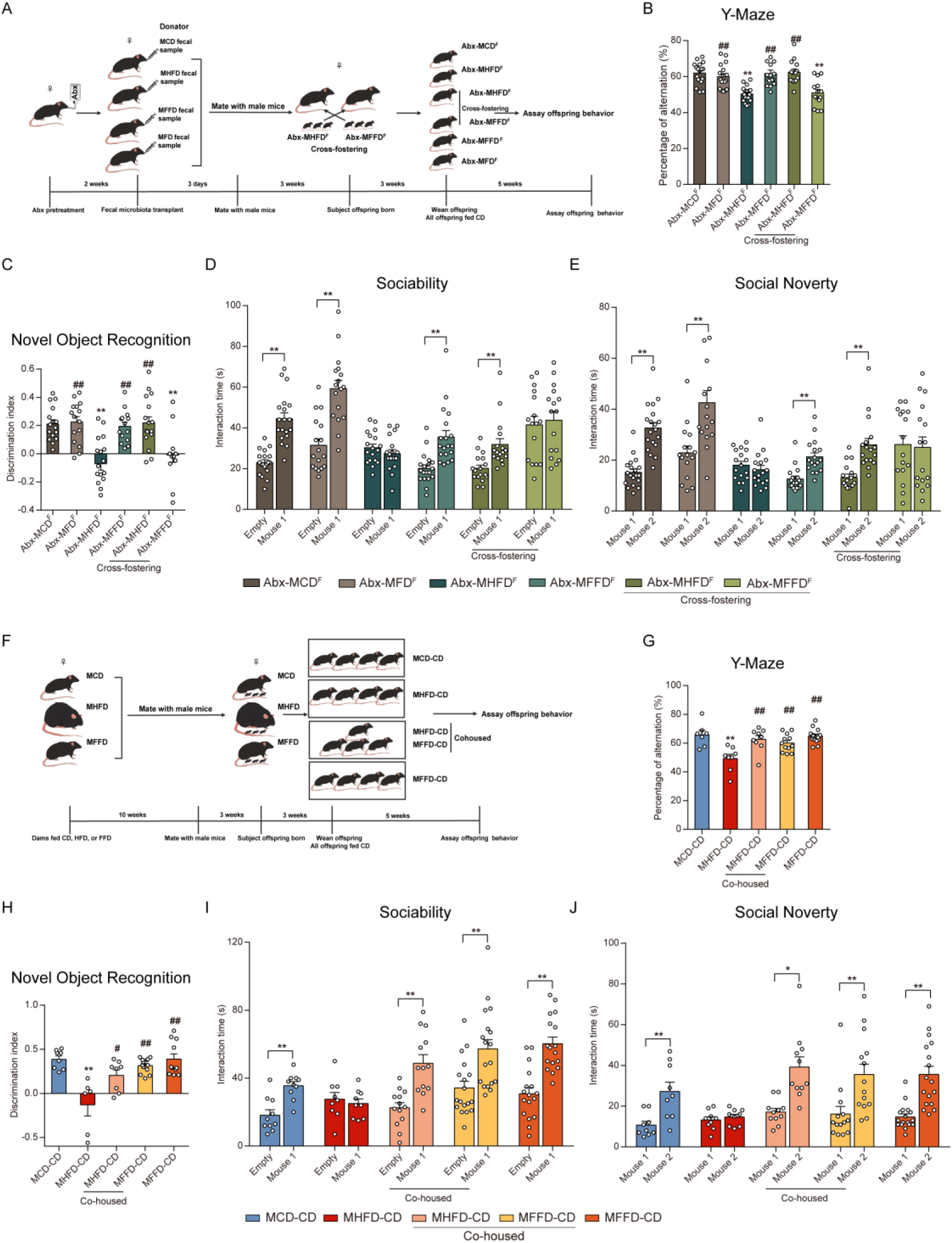
Gut Microbiota Mediates Maternal Obesity-Induced Cognitive and Social Deficits in Offspring. (A) Schematic of the maternal fecal microbiota transplant and crossing foster. (B-E) The offspring from fecal microbiota-transplanted mothers were tested by (B) Y-maze, (C) the novel object recognition test, (D) sociability, and (E) preference for social novelty (n = 13-17 mice/group). (F) Schematic of the offspring co-housing experiment. (G-J) The co-housed offspring were tested by (G)Y-maze, (H) the novel object recognition test), (I) sociability, and (J) preference for social novelty (n = 6-12 mice/group). Data presented as mean ± SEM. Statistical analyses were performed by one-way ANOVA with Tukey’s multiple comparison test (for B, C, G and H) or by paired two-tailed Student’s t test (for D, E, I, and J). For Tukey’s multiple comparison test, \**p* < 0.05, \*\**p* < 0.01, compared with the MCD-CD group, ^#^*p* < 0.05, ^##^*p* < 0.01, compared with the MHFD-CD group. For Student’s t test, \**p* < 0.05, \*\**p* < 0.01. See also Figure S5.

Subsequently, to investigate that maternal colonization with MHFD donor microbiota or MFFD donor microbiota influence behaviors in offspring after birth, cross-fostering experiments were performed by switching newborns between recipients of MHFD and MFFD’s microbiota. Strikingly, offspring derived from MHFD fecal microbiota-transplanted mothers, but reared by MFFD fecal microbiota-transplanted mothers, exhibited behavioral improvements; those from MFFD fecal microbiota-transplanted mothers that were reared by MHFD fecal microbiota-transplanted mothers exhibited behavioral impairments (**Figures 3A-3E, S5C and S5D**). Accordingly, MHFD offspring were co-housed with MFFD offspring, thus allowing the transfer of microbiota via coprophagy (**Figure 3F**). Similarly, MHFD offspring co-housed with MFFD offspring exhibited increased cognition, as well as improved sociability and preference for social novelty, as determined by behavioral tests (**Figures 3G-3J, S5E - S5G**). Altogether, these results indicate that there is a causality relationship between the gut microbiota disturbances and behavioral deficits in the offspring of maternal obesity, and maternal dietary fiber intake could protect against behavioral deficits of offspring via regulating gut microbiota.

### High Fiber Intake in Offspring’s Diet Restores Maternal Obesity-Induced Behavioral Deficits and Aberrant Spliceosome Alterations

We next sought to examine whether directly dietary fiber application to offspring could also reverse the behavioral and neurobiological deficits characteristic of MHFD offspring. MHFD offspring were fed FD from weaning (3 weeks) until adult, after which behaviors were tested (**Figures 4A, S6A and S6B**). Treatment with dietary fiber significantly improved cognition (*p* < 0.01), as well as sociability and the preference for social novelty (*p* < 0.05) (**Figures 4B-4E, S6C-S6D**).

**Figure 4.**
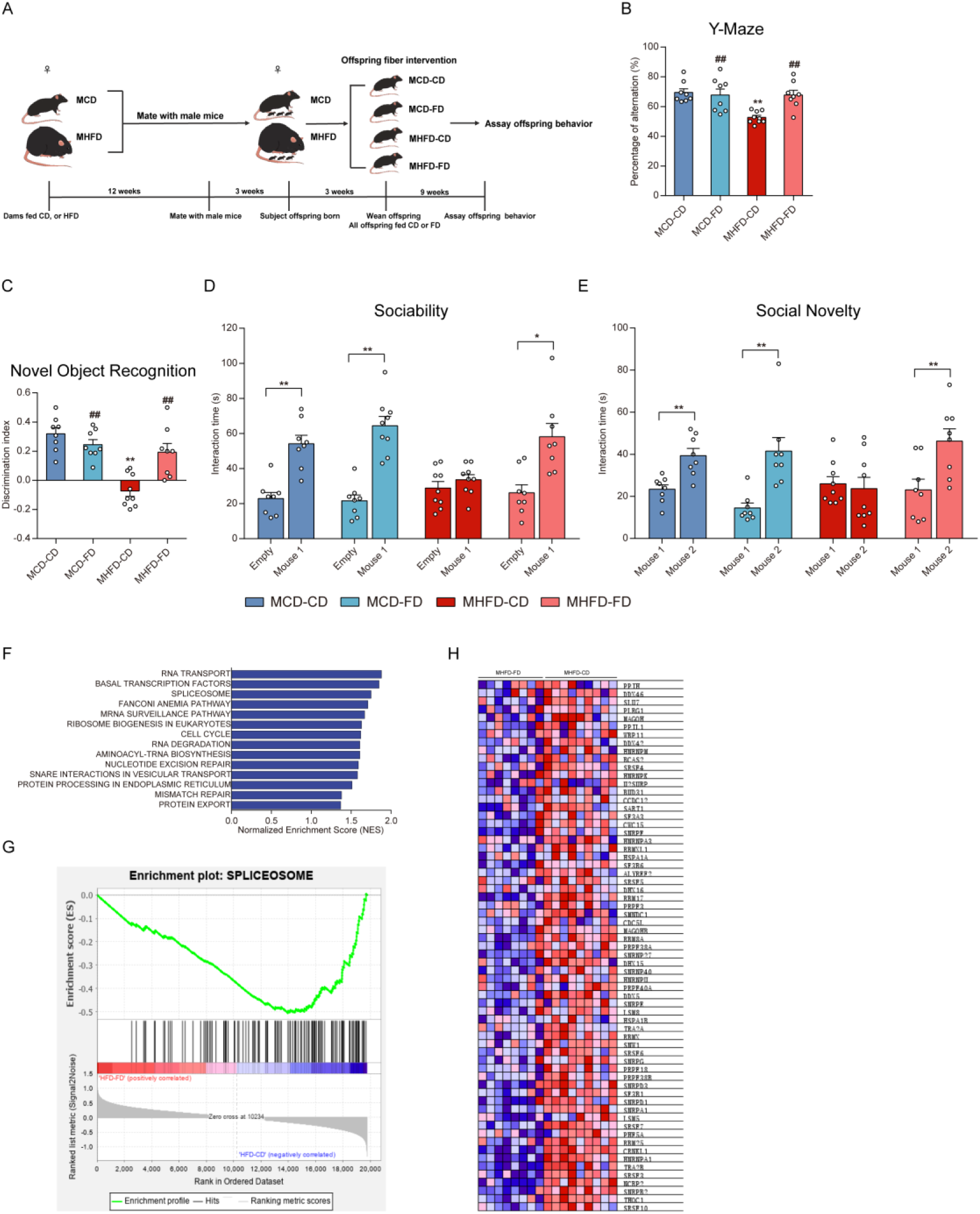
High Fiber Intake in Offspring’s Diet Restores Maternal Obesity-Induced Behavioral Deficits and Aberrant Spliceosome Alterations. (A) Schematic of the offspring directly dietary fiber administration. (B-E) Behavioral phenotypes in offspring after directly dietary fiber administration, including (B)Y-maze, (C) the novel object recognition test, (D) sociability, and (E) preference for social novelty (n = 8-9 mice/group). (F) KEGG pathways upregulated in the hippocampus of MHFD-CD mice by Gene set enrichment analysis (GSEA) (FDR < 25%) (n = 8-9 mice/group). (G-H) The enrichment plot (G) and the heat map of the leading-edge subset (H) for the spliceosome set. Data of (B) - (E) presented as mean ± SEM. Statistical analyses were performed by one-way ANOVA with Tukey’s multiple comparison test (for B and C) or by paired two-tailed Student’s t test (for D and E). For Tukey’s multiple comparison test, \**p* < 0.05, \*\**p* < 0.01, compared with the MCD-CD group, ^#^*p* < 0.05, ^##^*p* < 0.01, compared with the MHFD-CD group. For Student’s t test, \**p* < 0.05, \*\**p* < 0.01. See also Figure S6.

Gene expression profiling of RNA-seq identified 564 downregulated and 247 upregulated genes in the hippocampus of MHFD-FD and MCD-FD mice, using differentially expressed gene (DEG) analysis **(Figure S6E)** (FDR *q*-value<5%). There were relatively few sex-specific DEGs **(Figure S6F)**. Gene set enrichment analysis (GSEA) indicated that KEGG pathways involving transcription, translation, and protein quality control and export were upregulated in the hippocampus of MHFD-CD group mice, relative to MHFD-FD mice **(Figure 4F)**. A KEGG pathway involving RNA processing by the spliceosome was significantly upregulated in MHFD-CD mice (**Figures 4G** and **4H**) (FDR *q*-value < 25%). These data reveal that directly dietary fiber administration to 3 weeks old offspring could also effectively improve behavior and RNA splicing processes.

### High Fiber Intake in Offspring’s Diet Re-shapes the Gut Microbiome

We also examined whether dietary fiber could correct directly the changes in the microbiota of MHFD offspring. The gut microbiome from MHFD-FD offspring shifted away from that of MHFD-CD offspring **(Figures 5A and S6G**). Differentiating bacterial taxa belong predominantly to the Clostridiales and Bacteroidales orders, with single representatives from the Firmicutes phyla (**Figures 5B, S6H and S6I**). Consistent with maternal dietary fiber supplement, directly dietary fiber treatment improved the level of S24-7 (OTU_436_S24-7, OTU_516_S24-7 and OTU_573_S24-7 et al.) (**Figure 5C**). Specifically, *Bacteroides* (OTU_1_Bacteroides_acidifaciens, OTU_403_Bacteroidales, OTU_448_Bacteroides_acidifaciens, and OTU_672_Bacteroides_acidifaciens, et al.) were prevalent in all MHFD-FD samples and absent from most or all MHFD-CD samples (**Figure 5C**). Conversely, *Ruminococcus* (OTU_120_Ruminococcus, OTU_157_Ruminococcus and OTU_148_Ruminococcus_gnavus) was prevalent among most MHFD-CD mice and absent from MHFD-FD groups (**Figure 5C**). The abundance of the eight OTUs belonging to S24_7 positively correlated with increased long-term memory (R > 0.4, *p* < 0.05). Additionally, the level of *Bacteroides* significantly correlated with increased long-term memory and social behaviors (R > 0.5, *p* < 0.05). *Ruminococcus* showed the opposite effects, as it correlated with reduced memory behavior and social interaction deficits (R > 0.4, *p* < 0.05) (**Figure 5D**). These findings suggest that offspring dietary fiber intake could also correct the microbiota depletion in the offspring born to obese mothers.

**Figure 5.**
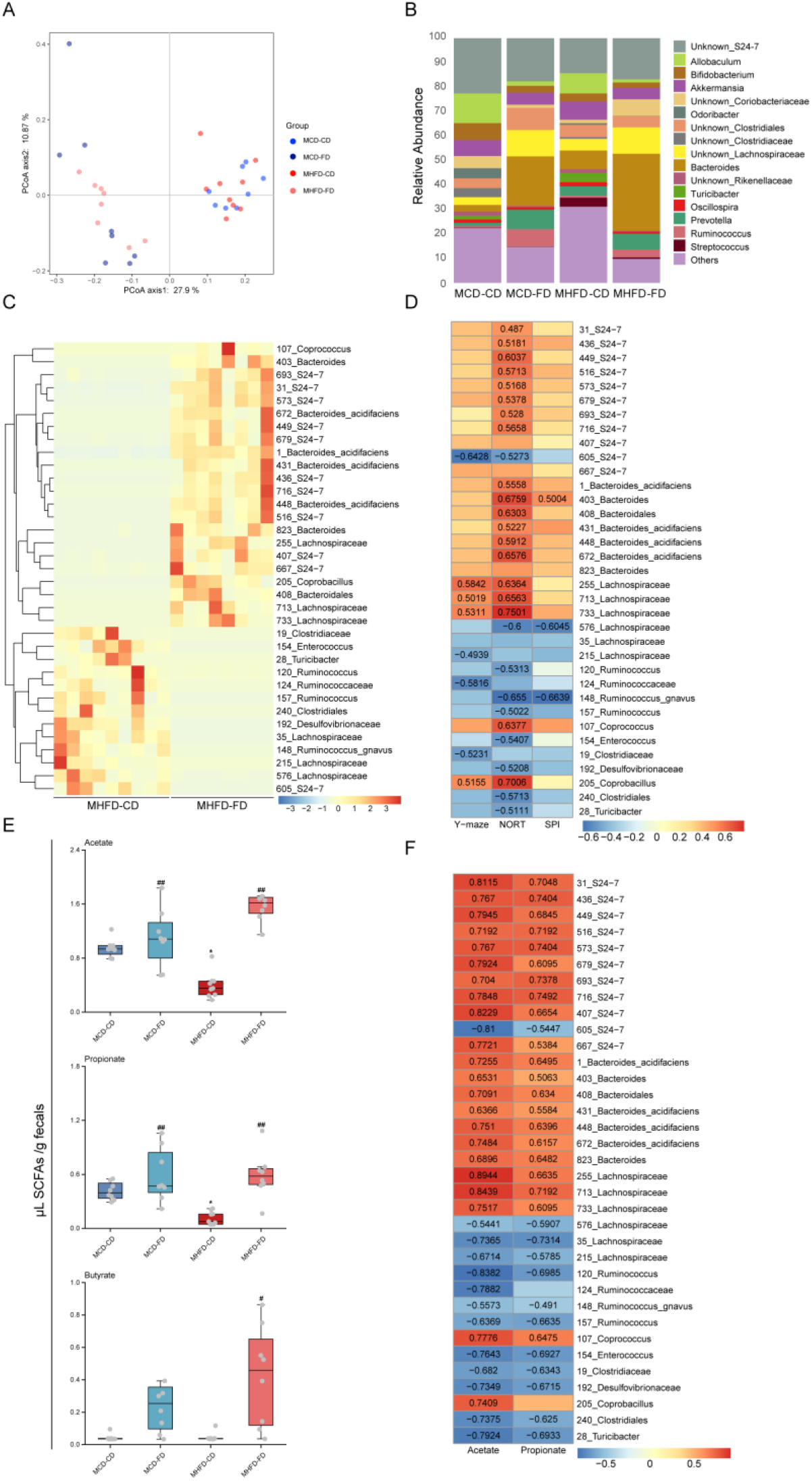
High Fiber Intake in Offspring’s Diet Re-shapes the Gut Microbiome. (A) UniFrac-based phylogenetic clustering(*p*<0.0001, R2=0.3395) (n = 8-9 mice/group). (B) The relative abundance of bacteria at the genus level. All genera with an average relative abundance below 1% were grouped to “others”. (C) A Z-score scaled heatmap of different OTUs identified by Wilcoxon rank-sum test between MHFD-CD and MHFD-FD with *p* ≤ 0.001. (D) Abundance of selected taxas in the offspring microbiome is correlated with behavior of offspring. Spearman’s rank correlation between the microbiome and mouse behaviors. If *p* value<0.05 for significant correlations, R is noted. (E) Short-chain fatty acid (SCFA) concentrations in feces (n = 8-9 mice/group). Data presented as median ± interquartile range. \**p* < 0.05, \*\**p* < 0.01, compared with the MCD-CD group, ^#^*p* < 0.05, ^##^*p* < 0.01, compared with the MHFD-CD group. Significant differences between mean values were determined by one-way ANOVA with Tukey’s multiple comparison test. (F) Spearman’s rank correlation between the selected taxas of microbiome and acetate/propionate. If *p* value<0.05 for significant correlations, R is noted. See also Figure S6.

### SCFAs Restores Maternal Obesity-Induced Cognitive and Social Behavioral Deficits in Offspring

In accord with maternal dietary fiber supplement, directly dietary fiber treatment elevated levels of acetate, propionate, but not butyrate in MHFD-FD offspring (**Figure 5E**). Furthermore, the level of acetate and propionate were significantly correlated with most differentiating bacterial taxa using Spearman’s rank correlation (**Figure 5F**) (R>0.4, *p* <0.05). We hypothesized that the selective decrease of acetate and propionate in the gut of MHFD offspring was causally related to their behavioral deficits. To test this hypothesis, acetate and propionate were introduced into the drinking water of MHFD offspring at weaning for 5 weeks, after which behaviors were tested (**Figures 6A and S7A**). Remarkably, treatment with a mix of acetate and propionate significantly improved cognition (*p* < 0.001), sociability and preference for social novelty in MHFD offspring (*p* < 0.001) (**Figures 6B-6E, S7B and S7C**). Accordingly, SCFAs treatment significantly improved the length and width of PSD (**Figures 6F and 6G**) (*p* < 0.01). Collectively, Together, these observations define the importance of SCFAs, which renders offspring resistant to social and cognitive impairments.

**Figure 6.**
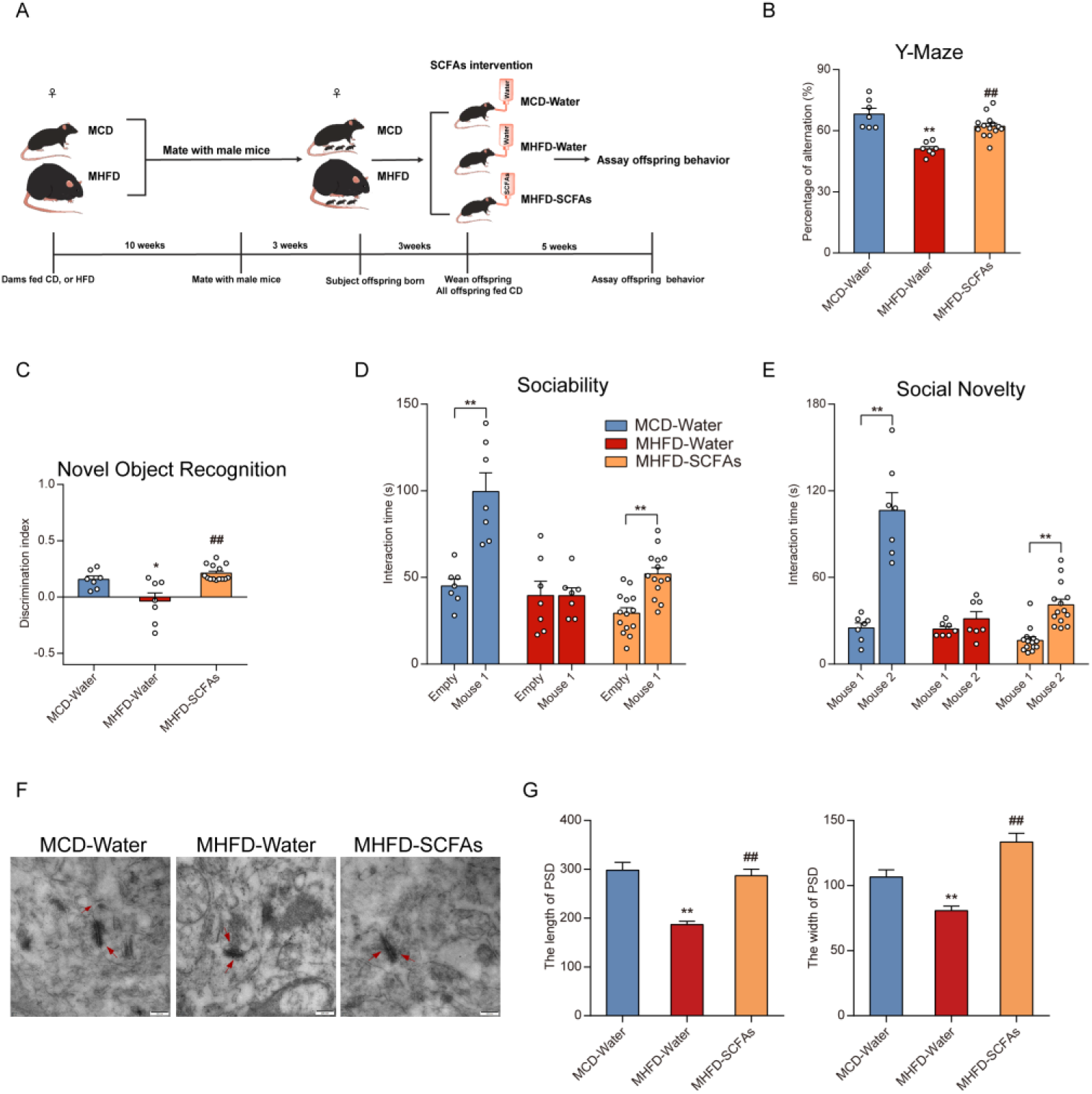
SCFAs Restores Maternal Obesity-Induced Cognitive and Social Behavioral Deficits in Offspring. (A) Schematic of treatment with a mixture of acetate and propionate. (B-E) Behavioral phenotypes in the offspring after treatment with a mixture of acetate and propionate, including (B) Y-maze, (C) the novel object recognition test, (D) sociability, and (E) preference for social novelty (n = 7-14 mice/group). (F) Representative images of ultrastructure of synapse. (G)The length (left) and width of PSD (right) (n = 3 mice/group). Data of (B) - (E) and (G) presented as mean ± SEM. Statistical analyses were performed by one-way ANOVA with Tukey’s multiple comparison test (for B, C and G) or by paired two-tailed Student’s t test (for D and E). For Tukey’s multiple comparison test, \**p* < 0.05, \*\**p* < 0.01, compared with the MCD-CD group, ^#^*p* < 0.05, ^##^*p* < 0.01, compared with the MHFD-CD group. For Student’s t test, \**p* < 0.05, \*\**p* < 0.01. See also Figure S7.

## DISCUSSION

In this study, we found that the maternal prepregnancy overweight and obesity are highly associated to the poorer cognitive performance and sociality of their children **(Table 1)**. Consistently, maternal obesity also induced cognitive and social behavioral deficits in the offspring of a mouse model **(Figure 1)**. Furthermore, high-fiber intake either in the maternal or in the offspring’s diet could improve the behavioral deficits, synaptic damage and aberrant spliceosome alterations in the offspring **(Figures 1, 2 and 4)**. Colonization of the gut microbiota from obese donors in the dams led to cognitive and social behaviors disorders in offspring. However, transplantation of the gut microbiota from high-fiber intake donors behaved better performance during the behavioral tests. These results indicated the causality relationship of the maternal gut microbiota composition disturbance and the behavioral changes in the offspring, which was further confirmed by the cross-fostering experiment **(Figure 3)**. Notably, the co-house experiment revealed that manipulation of the gut microbiome community of offspring after weaning could also alter the maternal obesity-induced behavioral deficits **(Figure 3)**. Moreover, SCFAs treatment in the offspring after weaning could also attenuate the maternal obesity-induced cognitive and social impairments **(Figure 6)**. These results further suggested that high fiber diet could protect against behavioral deficits of offspring possibly by restructuring the gut microbiome and enhancing the formation of the microbial metabolites.

Recent studies also reported that dietary fibers were able to improve cognitive performances and markedly modify behavior and brain chemistry relevant to anxiety and depression in rodent models (Burokas et al., 2017; Messaoudi et al., 2005). In the current study, it has been revealed that treatment of high fat diet-fed dams with dietary fiber could alleviate work and long-term memory impairments in both male and female offspring, which to the best of our knowledge, has not been reported before **(Figure 1)**. Of note, maternal dietary fiber intake could also restore the social behavior as well as preference for social novelty in both male and female offspring. Similarly, the high-dietary fiber intake in the offspring diet also mitigated the maternal obesity-induced cognitive and social deficits. RNA-seq analysis of the offspring’s hippocampi indicated that several disease-related pathways were regulated. Notably, offspring dietary fiber intake significantly reversed the upregulation of the KEGG pathway of spliceosome (mmu03040) in hippocampi **(Figure 4)**. Recent studies have highlighted the importance of aberrant alternative splicing of mRNA in the brains of subjects with an ASD diagnosis (Gandal et al., 2018; Gonatopoulos-Pournatzis et al., 2018). These discoveries provide experimental evidence that high dietary fiber intake can be an effective intervention to mitigate the influence of maternal obesity on the development of offspring’s central nervous system. Dietary fibers show broad impacts on various aspects of host physiology by adjusting the gut microbiota diversity. Here, we found that high fiber intake in the maternal det markedly altered the gut microbiota in the offspring of obese dams. Notably, OUTs belonging to S24_7 was found to be positively correlated with cognitive and social behaviors in mice **(Figure 2)**. However, to our best knowledge, the species of the S24_7 family microbes have not been fully understood. However, a recent study indicated that the microbes from the S24_7 family were highly related to the complex carbohydrate degradation in the gut (Lagkouvardos et al., 2019). Interestingly, Park et al. reported that the abundance of unclassified member of S24-7 was significantly decreased in APP/PS1 mice, a mouse model of Alzheimer’s disease (Park et al., 2017). Moreover, high-fiber intake in the offspring diet lowed the relative abundance of *Ruminococcus* and improved the *Bacteroides* **(Figure 5)**. The reasons why the influence of the high-fiber intake on the gut microbiota composition in the offspring diet is different from the maternal intervention could be related to the combination effects of mother-to-infant microbial transmission and shared environmental factors.

The possible explanation of how gut bacteria can affect the gene expression and behaviors host is by generating various metabolites (Hsiao et al.). SCFAs, including acetate and propionate are reported to be the key molecules modulating microglia maturation, morphology and function, so they could affect CNS function (Erny et al., 2015). Previous studies indicated that neurodevelopmental abnormality in ASD patients accompanied with impaired acetate and propionic acid metabolism (Liu et al., 2019; Wang et al., 2020). Consistently, it has been found that the concentrations of acetate and propionate in feces were reduced in offspring of obese dams. However, the formation of these SCFAs was restored by maternal or offspring dietary fiber intake, accompanied with the behavioral improvement **(Figures 2H and 5E)**. Furthermore, the SCFAs intervention experiments in the current research also indicated that SCFAs treatment significantly attenuated the maternal obesity-induced cognitive and social impairments in the offspring **(Figure 6)**. However, the precise mechanism by which SCFAs rescue social and cognitive behaviors remains to be determined. There is growing evidence indicates that acetate and propionate could regulate activity of histone deacetylases and promote acetylation of brain histones, which could be one of the future research targets to explain how these microbe metabolites improving the CNS function (Soliman and Rosenberger, 2011; Volmar and Wahlestedt, 2015; Waldecker et al., 2008). Taken together, our results demonstrate that the cognitive and social dysfunction associated with maternal obesity is induced by alterations of the offspring intestinal microbiota; and maternal or offspring dietary fiber intake can reverse behavioral dysfunction by regulating the bacterial composition and their metabolites SCFAs. This work provides new insight into the mechanism by which a marked shift in microbial ecology, caused by maternal high fat diet, can negatively impact both cognitive and social behaviors in offspring. These results may assist our deeper understanding the causation and underlying mechanisms linking maternal obesity with offspring neurodevelopment in human beings. Furthermore, we found that dietary fiber intervention for dams, or for 3-week old offspring from obesity mothers could both restore abnormal neurocognitive and social outcomes in offspring, which suggest that we have the opportunity to rectify some behavioral abnormalities of children during prenatal brain development, as well as postnatally. Overall, this finding opens new research avenues into preemptive therapies for neurodevelopmental disorders by targeting the maternal and/or offspring gut microbiota; and it also provides evidence that dietary fiber may be useful as a potential non-invasive, timely and tractable treatments for patients suffering from neurodevelopmental disorders.

## ACKNOWLEDGEMENTS

This work was financially supported by the National Key Research and Development Program of China (No. 2017YFD0400200), the National Natural Science Foundation of China (Nos. 81871118 and 81803231), the Innovative Talent Promotion Program-Technology Innovation Team (2019TD-006), the General Financial Grant from China Postdoctoral Science Foundation (No. 2016M602867), and the Special Financial Grant from China Postdoctoral Science Foundation (No. 2018T111104). Dr. Zhigang Liu is also funded by the Tang Cornell-China Scholars Program from Cornell University. We also thank Zixin He, Han Li, Junhe Zhao, and Hang Zhao (Northwest A&F University, China) for their help during animal experiments.

## AUTHOR CONTRIBUTIONS

X.L. (Xiaoning Liu), X.L(Xiang Li), B.X., X.J., Z.Z., S.Y. (Shikai Yan), L.L., S.Y. (Shufen Yuan), S.Z., X.D., M.H., Z.L., and X.L. (Xuebo Liu) performed the experiments and analyzed the data; X.L. (Xiaoning Liu), F.Y., E.C., R.L., B.Z., M.H., Z.L., and X.L. (Xuebo Liu) wrote the manuscript; X.L. (Xiaoning Liu), X.L(Xiang Li), X.J., Z.Z., S.Y. (Shikai Yan), M.H., Z.L., and X.L. (Xuebo Liu) prepared the figures. X.L. (Xiaoning Liu), H.M., Z.L., and X.L. (Xuebo Liu) supervised the project. All authors read and approved the final manuscript.

## DECLARATION OF INTERESTS

The authors declare no competing interests.

## EXPERIMENTAL MODEL AND SUBJECT DETAILS

### Human

Participants were 778 children (403 boys and 375 girls) aged 7-14 years (mean age =10.4±1.9) recruited from four primary schools in Taizhou and Wuxi. Sampling took place in classes randomly selected from each grade in the selected schools, where children and their parents were enrolled to complete a series of questionnaires. Those with a serious organic disease, abnormal physical development, or physical impairments were excluded from this study. The questionnaire included information on the child’s age, sex, body weight, height, physical activity, sleep, birth weight, breastfeeding history, maternal weight before pregnancy, weight gain during the period of pregnancy and height, parental education level, family income and a questionnaire of the Child Behavior Checklist (CBCL) (Shala and Dhamo, 2013). Anthropometric measures used standardized protocols (Lohman et al., 1988). Parents completed the social competence scale of the CBCL, which includes 20 social competence items with three social competence subscales measure competencies: activities (e.g., sports, hobbies); social subjects (e.g., friendships, interpersonal skills); and school performance (e.g., performance, ability, school problems) (Shala and Dhamo, 2013). The total social competence score is the sum of these three subscales. This study was approved by the Ethics Committee of School of Public Health of Shanghai Jiao Tong University for Human Subject Research, and all parents gave written informed consent, No. SJUPN-201815.

### Mice

Both C57Bl6/J male and female mice were obtained from Xi’an Jiaotong University (Xi’an, Shaanxi, China) at 7 weeks of age. Then mice were housed in the Northwest A&F University animal facility under standard conditions with a strict 12 h light/dark cycle (8:00 a.m.–8:00 p.m.), humidity at 50 ± 15%, temperature 22 ± 2 °C, and ad libitum access to food and water. Prior to the experimental period, all mice were maintained on chow, which we view as a reference diet. Littermates were randomly assigned to experimental groups. During the experiment, animal caretakers and investigators conducting the experiments were blinded to the allocation sequence. All of the experimental procedures were followed using the Guide for the Care and Use of Laboratory Animals: Eighth Edition (ISBN-10: 0-309-15396-4) and protocols were approved by the Northwest A&F University, and BGI Institutional Review Board on Bioethics and Biosafety (BGI-IRB).

### Maternal Dietary Fiber Treatment

During experiment, female mice were placed randomly on one of four diets: control diet (MCD), high fat diet (MHFD), a high fat diet with 37 g inulin per 1000 kcal as a source of fiber (MFFD), and a control diet with inulin as a source of fiber (MFD) for 10-12 weeks. The MCD and MFD diets consisted of 16.7% kcal from fat, 63.9% kcal from carbohydrates, and 19.4% kcal from protein. The MHFD and MFFD diets consisted of 60% kcal from fat, 20.6% kcal from carbohydrates, and 19.4% kcal from protein. The composition of all purified-ingredient diets used in this study are listed in Table S1. Female mice were five to a cage and weight was measured every six days before pregnancy. After 10 or 12 weeks on diets, females were paired with C57Bl6/J adult males fed chow to produce subject offspring (2 females: 1 male). Pregnant dams were single-housed and resulting offspring were weaned at 3 weeks of age. At weaning, different litters born within up to a week apart were combined and housed in a cage of 4-5 male or female mice per cage and all placed on CD, regardless of maternal diet (MCD, MHFD, MFFD, or MFD). All behavioral tests were performed in 8- to 10-week-old male and female mice. During subsequent analyses, investigators assessing, measuring or quantifying experimental outcomes were blinded to the intervention.

### Dietary Fiber Treatment to Offspring

Both female and male offspring from MHFD and MCD dams were switched to inulin containing diet (FD) at 3 weeks old, until 8- to 10-week-old age to be performed behavioral tests.

## METHOD DETAILS

### Microbiota depletion

In brief, mice were treated with an antibiotic solution (ATB) containing 0.5 g/L vancomycin, 1 g/L neomycin, 1 g/L ampicillin, and 1 g/L metronidazole added into the sterile drinking water. Solutions and bottles were changed once weekly respectively. Microbiota depletion was confirmed by qPCR. Mice received 14 days of ATB before undergoing fecal microbial transplantation (FMT).

### Gut microbiota transplantation

Microbiota transplantation was done according to a previous study (Wong et al., 2017). To transplant the ATB mice with microbiota, fecal samples were collected from four diet group donor mice. Fresh fecal samples were resuspended in sterile PBS (15 mL/g of feces) and then were vigorously vortexed for 5 min, followed by 5-min standing to precipitate particles. Supernatants were then used to colonize ATB mice by three consecutive days of oral gavage (200 μL/mouse). Colonized mice were subsequently mated with C57BL/6J male 3 days after colonization. And all pregnant mice were gavaged twice weekly during pregnancy and lactation. The colonization was confirmed by plating feces on BHI agar plates on anaerobic and aerobic conditions and by qPCR. For quantification of total bacterial burden, qPCR was performed with the universal bacteria-specific primer (8F: AGAGTTTGATCCTGGCTCAG, 338R: CTGCTGCCTCCCGTAGGAGT).

### Cross-fostering

Pregnant MHFD and MFFD mice were monitored every day, and immediately after birth (the day on which pups were born was considered P0. Pups were cross-fostered between P0 and P1.) all the MHFD pups were housed with lactating MFFD dams. Similarly, same day born MFFD pups were housed with lactating MHFD dams. After three weeks, the cross fostered pups were weaned and all placed on CD. All behavioral tests were performed in 8- to 10-week-old male and female mice.

### Co-housing

To co-housing experiments, sex-matched offspring from MHFD and MFFD dams were co-housed (3 weeks old) in gang cages at a ratio of 1:3 or 2:3 until 8- to 10-week-old age to be performed behavioral tests.

### SCFA Treatment

Both female and male offspring (3 weeks old) from MHFD dams treated with a mixture of sodium acetate (67.5 mM) and sodium propionate (22.5 mM) in drink water until 8- to 10-week-old age to be performed behavioral tests. Mice consumed the treated water ad libitum over the treatment period.

### Open Filed Testing

Mice were placed in an open arena (40 × 40 × 40 cm) and allowed to freely explore for 5 min while their position was continually monitored by an overhead camera and tracked using the SuperMaze software. Tracking allowed for measurement of distance traveled, speed, and position in the arena throughout the task. Time spent in the center of the arena, defined as the interior 20 × 20 cm, was recorded. Thoroughly clean the apparatus between mice using 70% ethanol.

### Novel Object Recognition

The object recognition test (ORT) is a relatively low-stress and efficient means for testing long-term learning and memory in mice (Lueptow, 2017). Because mice have an innate preference for novelty, if the mouse recognizes the familiar object, it will spend most of its time at the novel object. The ORT could be completed over 3 days: habituation day, training day, and testing day. On the habituation day, the mouse was removed from its home cage and placed in the middle of the open, empty arena (40 × 40 × 40 cm). Allow free exploration of the arena for 5 min. At the end of the session, the mouse was transferred to an empty holding cage. They were put back in home cage after all mice in the same home cage have been handled.

On the training day, two identical objects were placed in central symmetrical positions of the arena. Place the mouse in the center of the arena, equidistant from the two identical objects. After freely exploring the two objects for 5 min, the mouse was transported to the holding cage. On testing day, one of the training objects was replaced with a novel object. Then, the mouse was allowed to free explore for 5 min. Calculate the discrimination index (d2) as the time spent exploring the novel object minus the time spent exploring the familiar object divided by total exploration time.

### Y-Maze

The Y-maze task was performed as previous described (Arai et al., 2001) to assay the spontaneous alternation that was defined as successive entries into the three arms in overlapping triplet sets. The maze was made up of three arms, each of which was 35 cm long, 15 cm high and 5 cm wide, and converged to an equal angle (Shanghai Xinruan Information Technology, Shanghai, China). Mouse was placed in the center of the apparatus and was allowed to explore it for 8 min. The total numbers of arm entries and alternation were scored. The percent alternation was calculated as the ratio of actual to possible alternations (defined as the total number of arm entries −2) × 100%.

### Three-Chamber Social Test

The three-chamber test for sociability and preference for social novelty was performed as described (Buffington et al., 2016). Briefly, the mouse was first habituated to the full, empty arena for 5 min which was a 60 × 40 cm^2^ plexiglass box divided into three equally sized, interconnected chambers (left, center, right).

In the second 5-min, sociability was measured. During the period, the subject could interact either with an empty wire cup (empty) or an age and sex-matched stranger conspecific contained in the other wire cup (mouse 1). Time spent interacting (sniffing, crawling upon) with either the empty cup or the mouse 1 as well as time spent in each chamber were recorded using tracking software (SuperMaze), by independent observers.

Finally, preference for social novelty was assessed during a third 5-min period, by introducing a second stranger mouse (mouse 2) into the previously empty cup. Time spent interacting with either mouse 1 or mouse 2 as well as time spent in each chamber was recorded using the tracking software by independent observers.

### Mouse Fecal Sample Collection and 16S rRNA Microbiome Sequencing

Fecal samples were collected from respective groups at the before of the behavioral tests and stored in −80 °C until DNA extraction. The total cellular DNA was extracted with the E.Z.N.A. Stool DNA Kit (Omega, Norcross, GA, USA) according to the company instructions. Quantification of genomic DNA was verified using Qubit Fluorometer by using Qubit dsDNA BR Assay kit (Invitrogen, USA), and the quality was checked by running aliquot on 1% agarose gel. Qualified genomic DNA samples were processed for 16S rRNA library preparation as the method published previously. The bacterial hypervariable V3–V4 region of 16S rRNA was amplified by using primer: 341_F: 5′-ACTCCTACGGGAGGCAGCAG-3′ and 806_R: 5′-GGACTACHVGGGTWTCTAAT-3′. The validated library was used for sequencing on HiSeq2500 (Illumina, CA, USA) and generating 2×300 bp paired-end reads. The high quality paired-end reads were combined to tags based on overlaps by FLASH (Fast Length Adjustment of Short reads, v1.2.11), and then clustered into Operational Taxonomic Units (OTUs) at a similarity cutoff value of 97% using USEARCH (v7.0.1090), and chimeric sequences were compared with Gold database using UCHIME (v4.2.40) to detect. OUT representative sequences were taxonomically classified using Ribosomal Database Project (RDP) Classifer (v2.2) with a minimum confidence threshold of 0.6, followed by being mapped to the Greengenes database (v201305). All Tags back to OTU to get the OTU abundance statistics table of each sample using USEARCH_global. Principal coordinates analysis was performed by R (v3.5.2) at the OTU level to calculate unweighted and weighted Unifrac distance, followed by a permanova test (Vegan: Adonis) to detect differences among intervention groups. Alpha diversity was analyzed using R (v3.2.1) at the OTU level with Wilcoxon rank-sum test. Differential abundance analysis was performed using the Wilcoxon rank-sum test. The correlation was determined by R (v3.5.2) with Spearman’s rank correlations, followed by cor.test for multiple comparison corrections. *p* values<0.05 were considered significantly different.

### Short-Chain Fatty Acids Analysis

The concentrations of SCFAs (acetate, propionate, and butyrate) were determined using the gas chromatography. Approximately 150 mg of the fecal content sample was added in 1 mL water and vigorously vortexed. The samples were then mixed with 150 μL 50% H_2_SO_4_ and 1.6 mL diethyl ether. The suspensions were incubated on ice for 20 min to extract SCFAs, followed by centrifugation at 8000 rpm for 5 minutes at 4 °C. Lastly, the organic phase was collected and analyzed using a Shimadzu GC-2014C gas chromatograph (Shimadzu Corporation, Kyoto, Japan) equipped with a DB-FFAP capillary column (30 m × 0.25 μm × 0.25 mm) (Agilent Technologies, Wilmington, DE, USA) and flame ionization detector. The initial temperature was 50 °C, which was maintained for 3 min and then raised to 130 °C at 10 °C /min, increased to 170 °C at 5 °C /min, increased to 220 °C at 15 °C /min and held at this temperature for 3 min. The injector and the detector temperature were 250 °C and 270 °C, respectively.

### RNA Isolation and qRT-PCR

Total RNA was extracted from brain tissues using the RNAiso Plus (TaKaRa, Dalian, China). RNA concentrations were determined using NanoDrop 2000/2000C (Thermo Scientific; Waltham, MA, USA), and total RNA was reverse transcribed into cDNA using Evo M-MLV RT Kit with gDNA Clean for qPCR (Accurate Biotechnology (Hunan) Co., Ltd; Hunan, China), according to the manufacturer’s protocol using no more than 1 μg total RNA in 20 μL reactions. The mRNA expression was quantified using the SYBR green PCR kit (TaKaRa SYBRR Premix Ex Taq. II, Dalian, China) in a CFX96 Touch apparatus (Bio-Rad, Hercules, CA, USA) with the primers listed in KEY RESOURCES TABLE. The following conditions were used: 95 °C for 10 min and 95 °C for 15 s, followed by 40 cycles of 1 min at 60 °C. Difference in transcript levels were quantified by normalizing each amplicon to GAPDH, and the relative gene expression was calculated with the 2^−ΔΔCt^ method.

### RNA Sequence Analysis

Brain tissue from hippocampus was macro-dissected and stored in −80 °C until DNA extraction. Then the total RNA was extracted using TriZol (Invitrogen, Carlsbad, UA, USA) according to the manufacturer’s instructions, followed by being qualified and quantified using a NanoDrop and Agilent 2100 bioanalyzer (Thermo Fisher Scientific, MA, USA). RNA sequencing libraries were prepared using BGISEQ-500 (BGI-Shenzhen, China). Then the sequencing data were filtered and trimmed using Trimmomatic (v0.38) to obtain high-quality clean read data for sequence analysis. Clean reads were mapped to the Mus musculus genome sequence (ftp://ftp.ncbi.nlm.nih.gov/genomes/all/GCF/000/001/635/GCF_000001635.26_GRCm38.p6) using Hisat2 (v2-2.1.0). The reads of each sample were then assembled into transcripts and compared with reference gene models using StringTie (v1.3.4d). We merged the 49 transcripts to obtain a consensus transcript using a StringTie-Merge program. Transcripts that did not exist in the CDS database of the Mus musculus genome were extracted to predict new genes. The gene expression FPKM values were calculated using StringTie based on the consensus transcript. Differential expression analysis was performed using Ballgown (v2.12.0), an R programming-based tool designed to facilitate flexible differential expression analysis of RNA-Seq data. Only genes with FPKM >1 were subjected to analysis and the differential expression genes was determined (FDR-p < 0.05). After the gene expression FPKM values were calculated using StringTie software, gene-set enrichment analysis (GSEA) was performed on genes ranked by their differential expression Signal2Noise using GSEA software (v4.0.3). The KEGG pathway database was downloaded from https://www.kegg.jp/keggbin/search_pathway_text?map=mmu&keyword=&mode=1&viewImage=truehttps://www.kegg.jp/kegg. In this analysis, the normalized enrichment score (NES) and False Discovery Rate (FDR) were calculated for each gene set and compared among groups.

### Immunofluorescence

Immunofluorescence staining was performed according to previously described (Liu et al., 2017). The fixed brain sections were exposed to the primary antibodies at 4 °C overnight. After washing three times in PBS, these slices were incubated with a biotinylated goat anti-rabbit or goat anti-mouse at 37 °C for 20 min. The sections were then washed six times, followed by being mounted with mounting medium containing DAPI. Immunofluorescence images were acquired using an inverted fluorescent microscope (Olympus, Tokyo, Japan) (×200). Proteins were visualized using by Image J (developed by Wayne Rasband from NIH, USA) software.

### Transmission Electron Microscopy

A transmission electron microscope (TEM) analysis was done to assess the ultrastructure of synapse in the CA1 region of hippocampus as our previous research (Liu et al., 2020). In brief, CA1 region was split from hippocampus and treated in a cold fixative solution made of 2.5% glutaraldehyde (pH 7.2) at 4 °C for 24 h. After washing with PBS (0.1 M, pH 7.2), the specimens were post-fixed in 1% OsO4 (in 0.2M PBS, pH 7.2) at 4 °C for 1.5 h. After washing again with PBS, the specimens were dehydrated for 15 min in a series of ethanol solutions (30%, 50%, 70%, 80%, 90% (v/v), and then dehydrated in 100% (v/v) for 30 min twice. Samples were then infiltrated overnight in a mixture of LR-White resin (London Resin Company, Reading, U.K.) and alcohol (1:1, *v*/*v*), followed by infiltration with pure LR-White resin twice (for 6 h and 3 h, respectively) at room temperature. The samples were embedded in pure LR-White resin followed by being incubated at 60 °C for 48 h. Ultrathin sections were obtained with a diamond knife on the Leica EM UC7 ultramicrotome (Leica, Nussloch, Germany), and then stained with 3% aqueous solution of UA for 10-15 min, and re-floated by 4% Pb solution for 8-10 min. Sections were observed under JEM-1230 TEM (JEOL, Tokyo, Japan) at 80 kV, and were recorded with side-inserted BioScan Camera Model 792 (Gatan, Pleasanton, California, USA). The measurement was performed using Image J analysis software (National Institutes of Health, Scion Corporation, USA) by experimenters who blind to treatment group.

## QUANTIFICATION AND STATISTICAL ANALYSIS

To examine differences in the scores of the total social competence, activities, social and school between maternal weight status before the pregnancy, Kruskal-Wallis test was conducted. Associations of maternal prepregnancy BMI status with the participant characteristics were explored by using χ^2^ Tests and Kruskal-Wallis tests. Analyses were completed with the IBM SPSS program, version 22 (IBM, Chicago, IL, USA). The significance level was set to 0.05 for overall analyses.

RNA sequencing was analyzed using Ballgown (v2.12.0) and GSEA software (v4.0.3). Analysis and data visualization of microbial populations were carried out in R with various packages as described above. Other than RNA sequencing and gut microbiome statistical analysis was GraphPad Prism 7.0. ANOVA with Tukey’s post hoc analysis was used for parametric analysis of variance between groups, and two-tailed paired Student’s t test was used for pairwise comparisons, unless otherwise indicated. Data are presented as mean ± SEM and median ± interquartile range unless otherwise noted. *p* < 0.05 was considered significant. \**p* < 0.05, \*\**p* < 0.01, compared with MCD-CD group, ^#^*p* <0.05, ^##^*p* < 0.01 versus MHFD-CD group. The measurements were taken from distinct samples, no mice were excluded.

## Notes

### Competing Interest Statement

The authors have declared no competing interest.

### Summary of Updates

The figures were revised. Supplemental files updated.

