## Supplementary material for "Gut Microbiota Mediates High-Fiber Diet Alleviation of Maternal Obesity-Induced Cognitive and Social Deficits in Offspring": Methods & supplemental figures

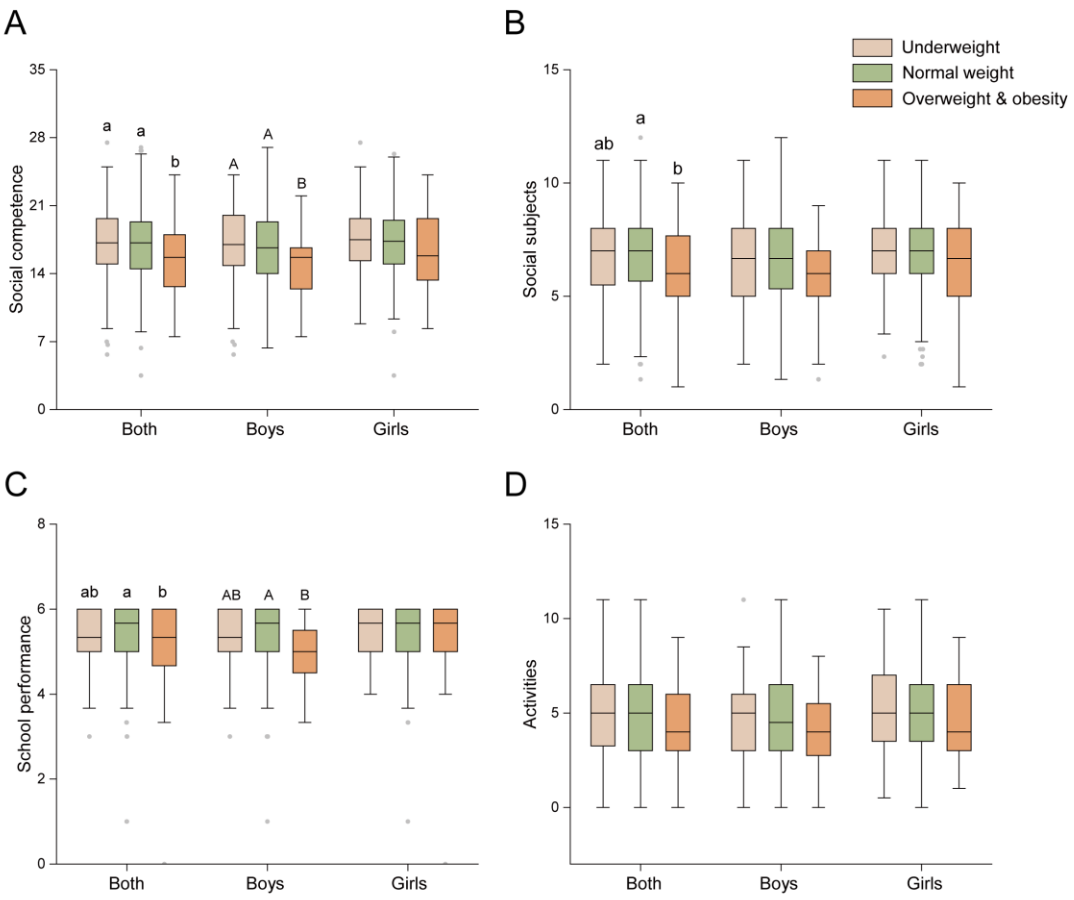


**Figure S1. Maternal Prepregnancy Overweight and Obesity are Associated with Impaired Child Neurodevelopment, Related to Table 1**

(A) Social competence of child.

(B) Social subjects of child.

(C) School performance of child.

(D) Activities of child.

Data presented as median ± interquartile range. Means with different letters (a, b, c) are significantly different from each other ($p < 0.05$). Significant differences between mean values were determined by Kruskal-Wallis test with Bonferroni's multiple comparisons.

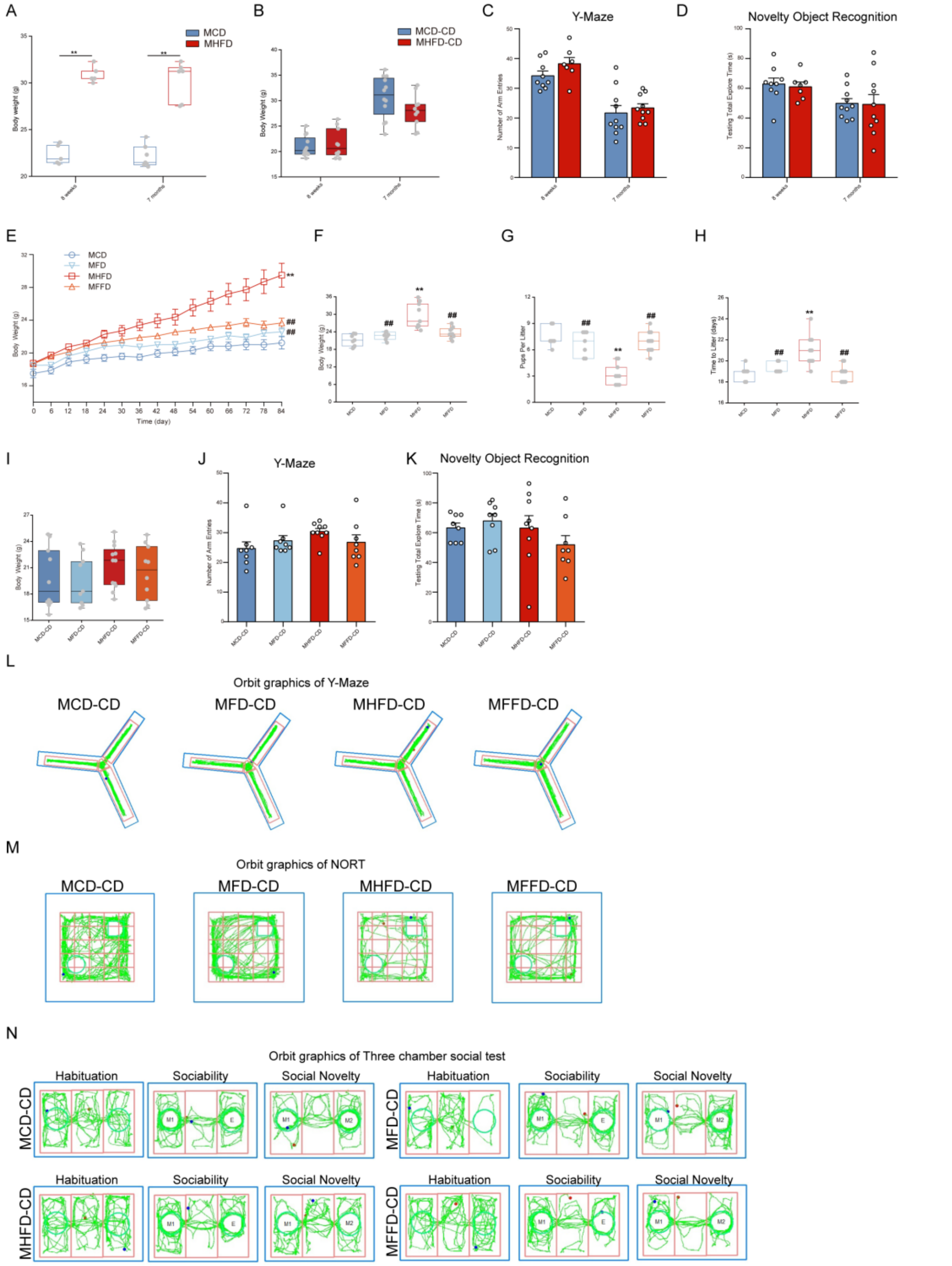

**Figure S2. Maternal Obesity and Dietary Fiber Intake Imprint the Weight of Mothers，Cognitive and Social Behaviors of Mouse Offspring, Related to Figure 1**

(A) Bodyweight of dams after 12 weeks on high fat diet (n = 5-7 mice/group).

(B) Offspring weight at either 8 weeks or 7 months of age (n =10-12 mice/group).

(C) Number of total arm entries in the Y-maze test (n = 7-10 mice/group).

(D) Testing total explore time in the novel object recognition test (n = 7-10 mice/group).

(E-F) Dam body weight after 12 weeks on diet (n = 7-10 mice/group).

(G-H) The litter size (G) and time to first litter (H) (n = 7-10 mice/group).

(I) Offspring weight at 8 weeks of age (n = 9-12 mice/group).

(J) Number of total arm entries (n = 8-9 mice/group).

(K) Testing total explore time (n= 8-9 mice/group).

(L-N) Representative exploratory activity of mice in the Y-maze test (L), in the novel object recognition test (M), and in the three-chamber test.

Data of (C) - (E) and (J) - (K) presented as mean ± SEM. Data of (A) – (B) and (F) – (I) presented as median ± interquartile range. For dams, $^{*}p < 0.05$, $^{**}p < 0.01$, compared with MCD group, $^{\#}p < 0.05$, $^{\#\#}p < 0.01$ versus MHFD group. For offspring, $^{*}p < 0.05$, $^{**}p < 0.01$, compared with MCD-CD group, $^{\#}p < 0.05$, $^{\#\#}p < 0.01$ versus MHFD-CD group. Significant differences between mean values were determined by one-way ANOVA with Tukey’s multiple comparisons test.

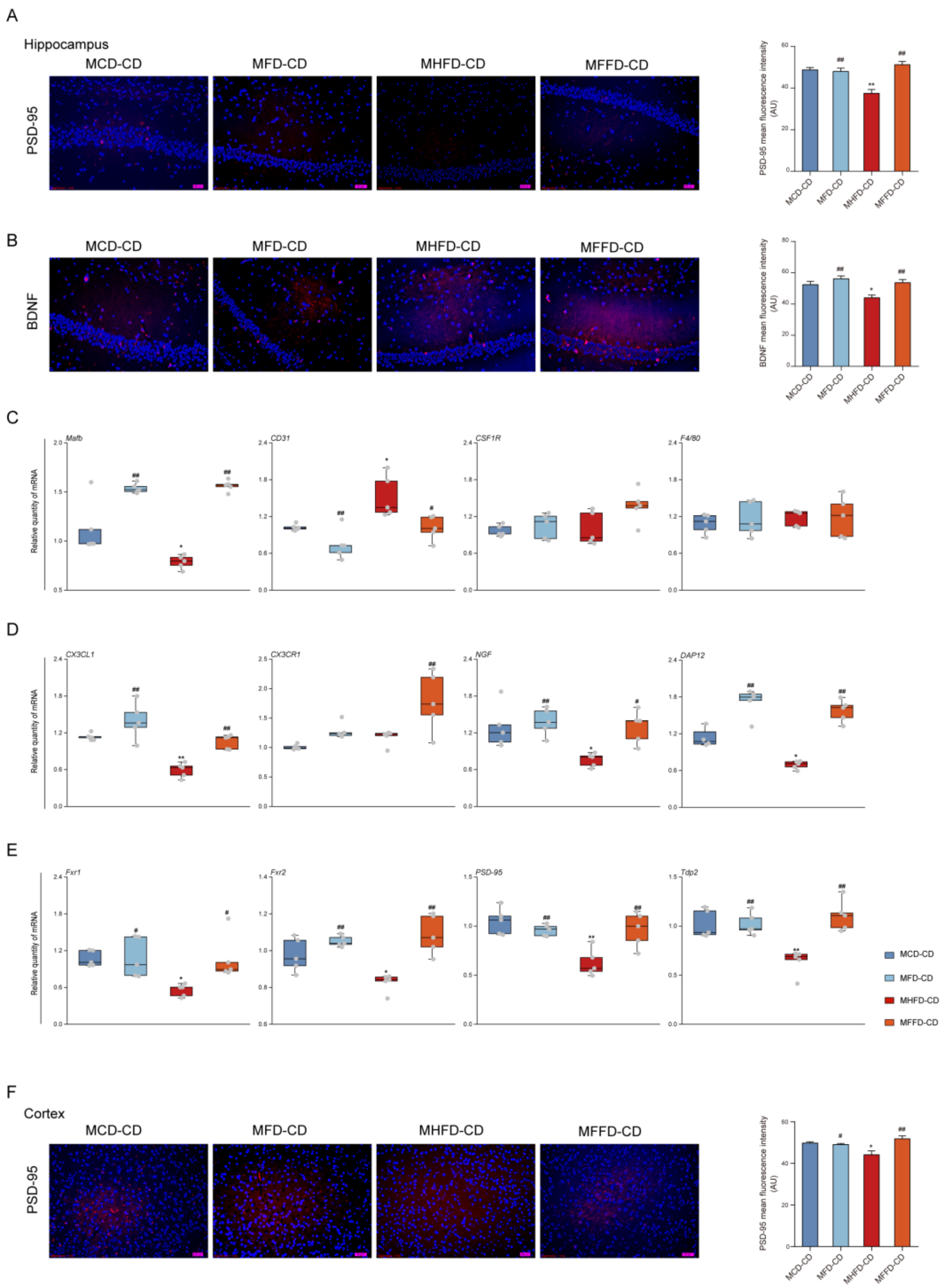


**Figure S3. Dietary Fiber Intake Restores Maternal Obesity-Induced Synaptic Damage and Disruption of Microglia Maturation in Offspring，Related to Figure 2**

(A) Representative immunofluorescence images (left) and immunofluorescence intensity (right) of PSD-95 in the hippocampus (n = 6 mice/group).

(B) Representative immunofluorescence images (left) and immunofluorescence intensity (right) of BDNF in the hippocampus (n = 6 mice/group).

(C) The mRNA expressions of *Mafb*, *CD31*, *CSF1R* and *F4/80* in the prefrontal cortex (n = 5 mice/group).

(D) The mRNA expressions of *CX3CL1*, *CX3CR1*, *NGF* and *DAP12* in the prefrontal cortex (n = 5 mice/group).

(E) The mRNA expressions of *Fxr1*, *Fxr2*, *PSD-95* and *Tdp2* in the prefrontal cortex (n = 5 mice/group).

(F) Representative immunofluorescence images (left) and immunofluorescence intensity (right) of PSD-95 in the prefrontal cortex (n = 6 mice/group).

Data of (A) - (B) and (F) presented as mean ± SEM. Data of (C) – (E) presented as median ± interquartile range. $^{*}p < 0.05$, $^{**}p < 0.01$, compared with MCD-CD group, $^{\#}p < 0.05$, $^{\#\#}p < 0.01$ versus MHFD-CD group. Significant differences between mean values were determined by one-way ANOVA with Tukey's multiple comparisons test.

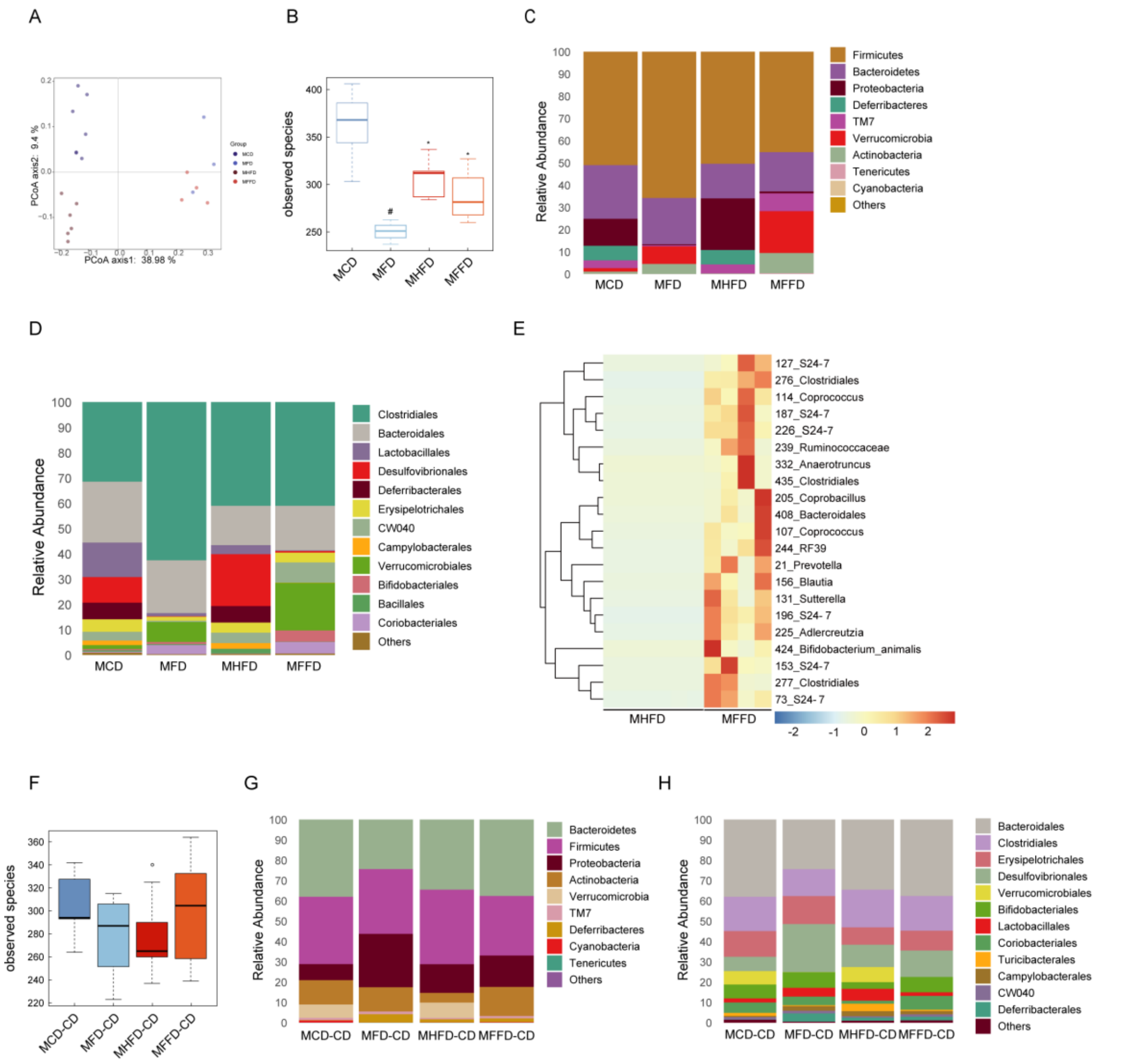


**Figure S4. Dietary Fiber Intake Re-Shapes the Gut Microbiome in Both Mother and Offspring Mice, Related to Figure 2**

(A) Principal coordinate analysis (PCoA) based on unweighted Unifrac distance and permanova (adonis) were used to test the difference in gut microbiota composition and diversity between groups of dams ($p < 0.001$, $R^2 = 0.52$) (n = 3-7 mice/group).

(B) α-Diversity in dams as measured by observed operational taxonomic units (OTUs) from 16S rRNA gene sequencing. Differences between treatment groups were tested by Wilcoxon rank-sum test ($^{*}p < 0.05$).

(C-D) The relative abundance of bacteria (C) at the phylum level and (D) at the order level in dams.

(E) A Z-score scaled heatmap of different OTUs identified by Wilcoxon rank-sum test between MHFD and MFFD with $p \leq 0.01$.

(F) α-Diversity in offspring as measured by observed operational taxonomic units (OTUs). Differences between treatment groups were tested by Wilcoxon rank-sum test (n = 8-9 mice/group).

(G-H) The relative abundance of bacteria (G) at the phylum level and (H) at the order level in offspring.

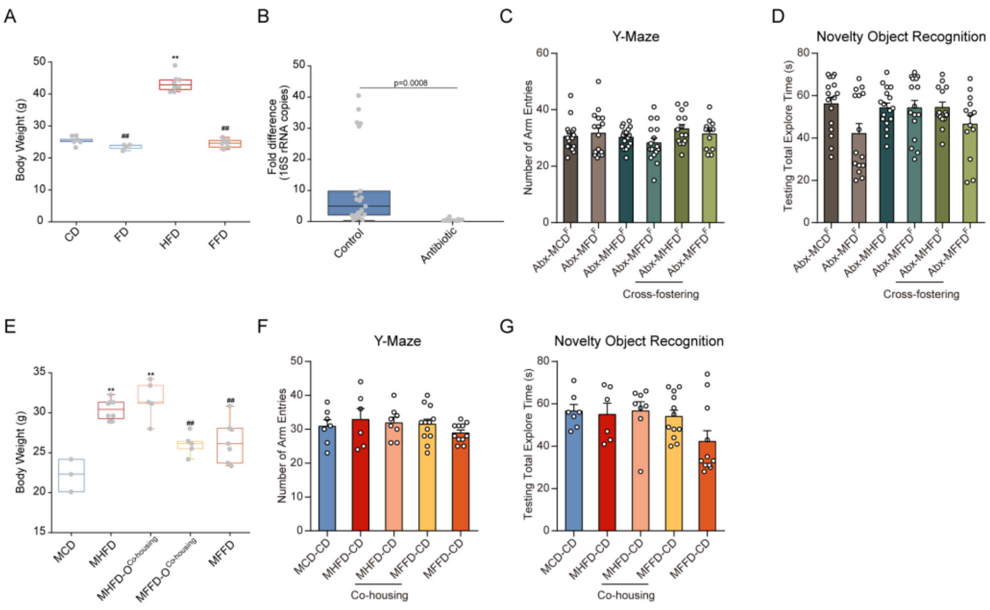


**Figure S5. Gut Microbiota Mediates Maternal Obesity-Induced Cognitive and Social Deficits in Offspring, Related to Figure 3**

(A) Body weight of female mice donor fed one of four diets (n = 4-8 mice /group).

(B) qPCR of 16S rDNA analysis to ensure the removal efficacy of microbiota (n = 20-21 mice /group).

(C-D) Number of total arm entries of offspring from fecal microbiota-transplanted mothers in the Y-maze test (n = 13-17 mice/group).

(D) Testing total explore time of offspring from fecal microbiota-transplanted mothers in the novel object recognition test.

(E) Body weight of dams, who gave birth to co-housed offspring (n = 3-8 mice/group).

(F) Number of total arm entries of co-housed offspring in the Y-maze test (n = 6-12 mice/group).

(G) Testing total explore time of co-housed offspring in the novel object recognition test (n = 6-12 mice/group).

Data of (C) - (D) and (F) –(G) presented as mean ± SEM. Data of (A) – (B) and (E) presented as median ± interquartile range. For donor mice, $^{*}p < 0.05$, $^{**}p < 0.01$, compared with CD group, $^{\#}p < 0.05$, $^{\#\#}p < 0.01$ versus HFD group. For dams, $^{*}p < 0.05$, $^{**}p < 0.01$, compared with MCD group, $^{\#}p < 0.05$, $^{\#\#}p < 0.01$ versus MHFD group. For offspring, $^{*}p < 0.05$, $^{**}p<0.01$, compared with MCD-CD group, $^{\#}p < 0.05$, $^{\#\#}p < 0.01$ versus MHFD-CD group. Significant differences between mean values were determined by one-way ANOVA with Tukey's multiple comparisons test.

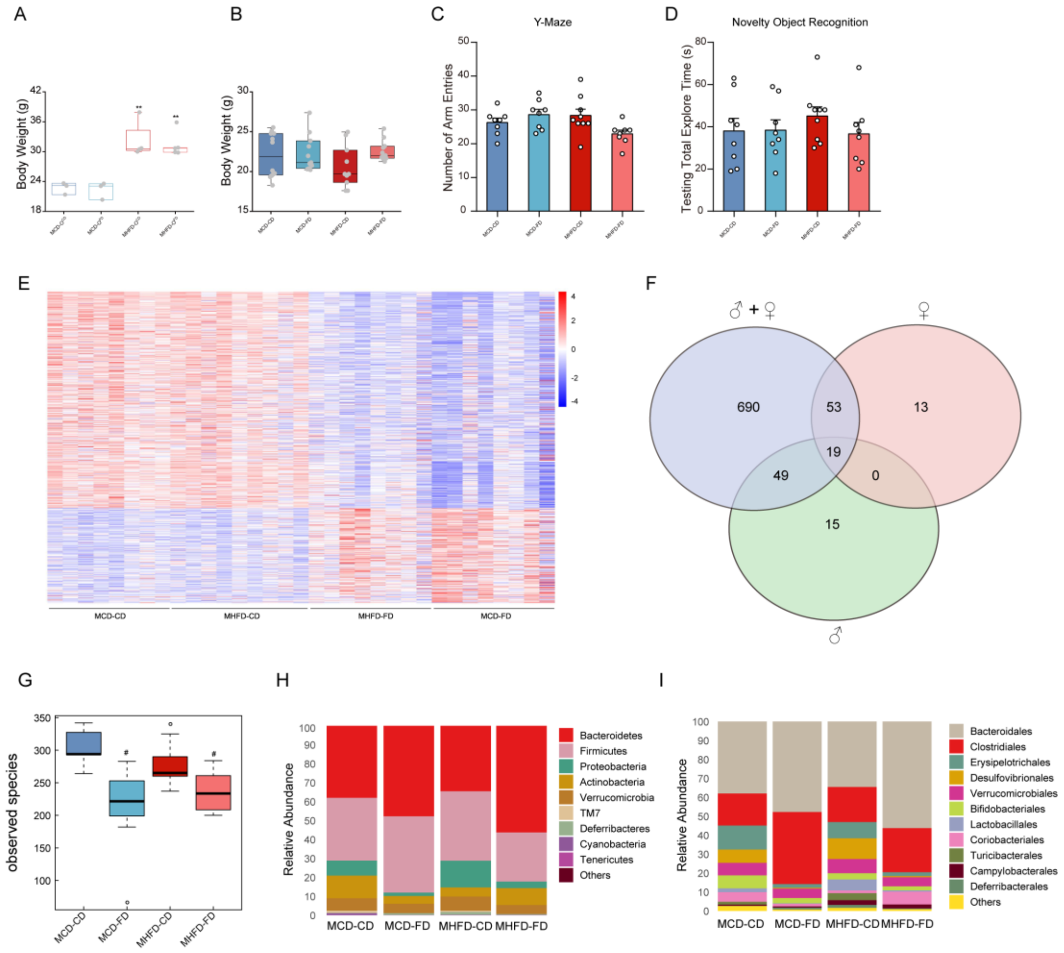

**Figure S6. Offspring Dietary Fiber Intake Restores Maternal Obesity-Induced Gene Changes in the Hippocampus and Microbiota Depletion, Related to Figure 4 and Figure 5**

(A) Body weight of dams (n = 3-5 mice/group).

(B) Body weight of offspring (n = 10-12 mice/group).

(C) Number of total arm entries of offspring after directly dietary fiber administration in the Y-maze test (n = 8-9 mice/group).

(D) Testing total explore time of offspring after directly dietary fiber administration in the novel object recognition test (n = 8-9 mice/group).

(E) Heatmap of the DEGs of the hippocampus using Ballgown software (FDR-$p < 0.05$).

(F) Overlap of the DEGs between gender combinations (n = 8-9 mice/group).

(G) α-Diversity in offspring as measured by observed operational taxonomic units (OTUs). Differences between treatment groups were tested by Wilcoxon rank-sum test (n = 8-9 mice/group).

(H-I) The relative abundance of bacteria (H) at the phylum level and (I) at the order level in offspring.

Data of (C) - (D) presented as mean ± SEM. Data of (A) – (B) presented as median ± interquartile range. For dams, $^{*}p < 0.05$, $^{**}p < 0.01$, compared with MCD group, $^{\#}p < 0.05$, $^{\#\#}p < 0.01$ versus MHFD group. For offspring, $^{*}p < 0.05$, $^{**}p<0.01$, compared with MCD-CD group, $^{\#}p < 0.05$, $^{\#\#}p < 0.01$ versus MHFD-CD group. Significant differences between mean values were determined by one-way ANOVA with Tukey's multiple comparisons test.

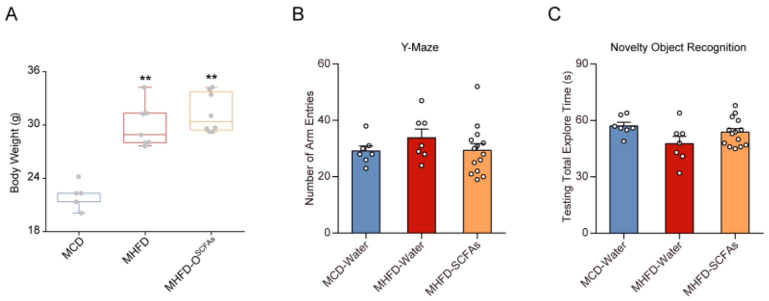


**Figure S7. SCFAs Supplementation Restores Maternal Obesity-Induced Cognitive Behavior Deficits in Offspring, Related to Figure 6**

(A) Body weight of dams (n = 5-8 mice/group).

(B) Number of total arm entries of offspring in the Y-maze test (n = 7-14 mice/group).

(C) Testing total explore time of offspring in the novel object recognition test (n = 7-14 mice/group).

Data of (B) - (C) presented as mean ± SEM. Data of (A) presented as median ± interquartile range. For dams, $^{*}p < 0.05$, $^{**}p < 0.01$, compared with MCD group, $^{\#}p < 0.05$, $^{\#\#}p < 0.01$ versus MHFD group. For offspring, $^{*}p < 0.05$, $^{**}p<0.01$, compared with MCD-CD group, $^{\#}p < 0.05$, $^{\#\#}p < 0.01$ versus MHFD-CD group. Significant differences between mean values were determined by one-way ANOVA with Tukey's multiple comparisons test.
